# Wide-Field Calcium Imaging of Dynamic Cortical Networks During Locomotion

**DOI:** 10.1101/2020.07.06.189670

**Authors:** Sarah L. West, Justin D. Aronson, Laurentiu S. Popa, Russell E. Carter, Kathryn D. Feller, William M. Chiesl, Morgan L. Gerhart, Aditya C. Shekhar, Leila Ghanbari, Suhasa B. Kodandaramaiah, Timothy J. Ebner

## Abstract

Motor behavior results in widespread activation of the cerebral cortex. Therefore, fully understanding the cerebral cortex’s role in motor behavior requires a mesoscopic level description of the engaged cortical regions and their functional interactions. Mesoscopic imaging of Ca^2+^ fluorescence through transparent polymer skulls implanted on transgenic Thy1-GCaMP6f mice reveals widespread activation of the cerebral cortex during locomotion, including not only in primary motor and somatosensory regions but also in secondary motor, retrosplenial, and visual cortices. Using the time series of Ca^2+^ fluorescence from 28 regions (nodes) we found to be consistent across mice, we examined the changes in functional connectivity from rest to locomotion. At the initiation of locomotion, the most anterior nodes of the secondary motor cortex increase in correlation with other nodes, while other nodes decrease in correlation. Eigenvector centrality confirms these changes in functional connectivity. Directed Granger causality analysis reveals an increase in causal influence of anterior regions of secondary motor cortex on other dorsal cortical regions at the onset of locomotion. These results highlight the global changes in correlation, centrality, and causality occurring throughout the cerebral cortex between rest and locomotion and suggest that the premotor areas play an important role in organizing these changes.

---

Behavior involves processing and integrating information within and across multiple brain regions, and a prominent view is that sensory, motor, and cognitive behaviors emerge from the neuronal interactions in and among these regions (Harris 2005; Buzsaki 2010; Carrillo-Reid et al. 2016; Allen et al. 2017; Musall et al. 2019; Steinmetz et al. 2019). Even simple sensory stimuli and motor tasks involve multiple cortical areas; for example, deflection of a single whisker results in activation across the sensorimotor cortices (Ferezou et al. 2007), while decision making relies on sensory signals that widely engage the cortex (Hernandez et al. 2010; Makino et al. 2017; Allen et al. 2017; Pinto et al. 2019). Notably, movements cause widespread activation across the cerebral cortex, even in regions not directly involved in motor control (Steinmetz et al. 2019; Musall et al. 2019), and this activity contains mixed representations of sensory, motor, and behavioral state (Li 2015; Kauvar et al. 2020). This distributed processing has been observed in a wide range of contexts, including in instructed and uninstructed discrete movements, in locomotion, and across measurement modalities (Makino et al. 2017; Clancy et al. 2019; Musall et al. 2019; Kauvar et al. 2020). The cerebral cortex translates sensory information into accurate motor plans (Freedman and Ibos 2018; Serino 2019) and uses motor information to accurately interpret sensory stimuli (for review see (Schneider 2020). Widespread projections from pyramidal neurons to different functional regions provide the anatomical substrate for these global computations (Zingg et al. 2014; Economo et al. 2018; Han et al. 2018). Therefore, the cerebral cortex’s role in a motor behavior cannot be easily understood by investigating its components in isolation, emphasizing the need to study many cortical regions simultaneously to determine how they coordinate and integrate activity to produce a unitary behavior.

During movement, various regions of the cerebral cortex are thought to be tasked with the integration of sensory information used for motor planning. The posterior parietal cortex (PPC; retrosplenial cortex in rodents) is involved in integrating sensory information of multiple modalities and with navigating in space (Mao et al. 2020). During locomotion in the cat, the PPC is necessary for storing information of an upcoming obstacle (Drew and Marigold 2015; Takakusaki 2017) and in the mouse, the retrosplenial cortex shows increased correlation with sensory cortical regions (Clancy et al. 2019). In non-human primates, multimodal neurons in the premotor cortex encode spatial information of visual, auditory, and somatosensory stimuli that occur close to the body. The premotor cortex uses these information to plan movement trajectories that are executed by the primary motor cortex (M1)(Di Pellegrino G. and Ladavas 2015). The major upstream premotor cortical region in rodents, the secondary motor cortex (M2), is thought to perform functions analogous to those performed by primate premotor and supplementary motor cortices (Inagaki et al. 2018; Yang and Kwan 2020). M2 encompasses more rostral regions, such as the anterior lateral motor area, the rostral forelimb area, and medial secondary motor cortex (Neafsey and Sievert 1982; Hira et al. 2013; Inagaki et al. 2018; Yang and Kwan 2020).

In return, movement alters neuronal activity in primary sensory regions. Neuronal firing in the primary visual cortex (V1), both spontaneous and in response to stimuli, increases during movements (Niell and Stryker 2010; Saleem et al. 2013; Lee et al. 2014; Dadarlat and Stryker 2017; Dipoppa et al. 2018), while neural activity decreases in the primary auditory cortex (Schneider et al. 2014; Zhou et al. 2014; McGinley et al. 2015; Schneider and Mooney 2018). Spontaneous activity in somatosensory cortex also increases (Chapin and Woodward 1982); (Favorov et al. 2015; Ayaz et al. 2019). Notably, these modulations in neural firing occur even in the absence of a change in sensory input, suggesting the changes are internally driven by the motor state (see (Schneider 2020)). Since performance on visual and auditory discrimination tasks increases and decreases, respectively, during locomotion (Schneider and Mooney 2018; Tang and Higley 2020), these changes likely reflect re-prioritizing incoming stimuli and sensory modes to provide the most useful information during movement (Schneider 2020; Parker et al. 2020). Supporting this concept, pyramidal neurons in the primary visual cortex activated during locomotion are strongly driven by unexpected stimuli (Keller et al. 2012).

Because these changes in sensory encoding are due to motor activity, information flows not only from sensory to other regions, but feed-forward signals flow from motor to sensory regions. However, the mechanism of this motor-driven sensory modulation is unclear. While cholinergic inputs from the mesencephalic locomotor region (MLR) drive modulation in V1 neurons similar to that found during locomotion (Lee et al. 2014), sensory modulation in visual and auditory cortices occurs most strongly in pyramidal neurons of layers II/III, suggesting a cortical origin of motor-driven changes (Polack et al. 2013; Zhou et al. 2014). There is evidence that these signals originate from primary and premotor cortical regions and act as an “efference copy” of ongoing motor activity (Lee et al. 2013; Zagha et al. 2013; Schneider et al. 2014; Nelson and Mooney 2016; Leinweber et al. 2017).

Primary motor cortex plays a crucial role in the accurate placement of limbs during locomotion. While much of the locomotor limb pattern is produced by central pattern generators in the spinal cord (Josset et al. 2018; Sharma et al. 2019), lesioning M1 or the corticospinal tract produces locomotor deficits, including hypermetria and abnormalities in limb trajectory and intralimb coordination (Liddell and Phillips 1944). Pyramidal neuronal firing in M1 correlates with individual muscle activity during locomotion (Drew et al. 2008a; Drew et al. 2008b; Drew and Marigold 2015) and modulates when the subject maneuvers around an obstacle. Less is known about the function of the premotor cortex in locomotion (Drew and Marigold 2015), but M2 in rodents develops causal influence over activity in much of the cortex after a mouse learns a discrete motor task (Makino et al. 2017).

The functional interactions among motor, premotor, sensory, and associative cortices must be investigated simultaneously to understand how the exchange of sensory and motor information is organized across the cerebral cortex during motor behaviors. This approach can provide insights into the function of individual regions, as well as elucidate the functional dynamics of the cerebral cortex as a unit. Here, we examine the interactions among different cortical areas during the transition from awake quiescence (rest) to continued, steady-state locomotion using mesoscopic Ca^2+^ imaging across the entire dorsal cerebral cortex in head-fixed mice. The increases in Ca^2+^ fluorescence correlate with the parameters of locomotion. Analysis of the fluorescence activity between cortical regions at different time periods, from rest to locomotion and back to rest, reveal an evolution in the patterns of functional connectivity, as measured by correlation, centrality, and Granger causality. Most notably, primary motor, primary sensory, and parietal cortical regions exhibit decreases in functional connectivity, while retrosplenial and the anterior regions in M2 exhibit increases. Moreover, the anterior regions in M2 alone show increases in outward Granger causality to many other regions, indicating they may play an important role in coordinating motor and sensory computations across the cerebral cortex during locomotion.

## MATERIALS AND METHODS

All animal studies were approved by and conducted in conformity with the Institutional Animal Care and Use Committee of the University of Minnesota.

### Animals and surgical procedures

Eight (5 male, 3 female) transgenic mice expressing GCaMP6f primarily in excitatory pyramidal neurons of the cerebral cortex (C57BL/6J, Thy1-GCaMP6f Jackson laboratories JAX 024339) were used (Dana et al. 2014). To obtain optical access to a large region of the dorsal cerebral cortex, we implanted morphologically conformant windows made from transparent polymer (Ghanbari et al. 2019). Prior to surgery, animals were administered slow release buprenorphine (2mg/kg s.c.) and then anesthetized with isoflurane (5% induction, 0.5-3% maintenance). The head was shaved and mounted in a stereotaxic frame that did not damage the auditory meatus. Depth of anesthesia was monitored by toe pinch response every 15 min. Isoflurane levels were adjusted with respiration rate or if a response to pain was registered. Body temperature was maintained (37°C) using a feedback-controlled heating pad, and the corneas were protected with eye ointment. The surgical procedure began with excision of the scalp, followed by removal of the facia so that the positions of lambda and bregma could be recorded. High resolution images with a reference scale were captured both before and after securing the implant to the skull using a digital microscope camera (S01-0801A, Science Supply) attached to the surgical microscope to identify bregma after removing the skull. A manual craniotomy removed a flap of skull that matched the geometry of the implant window, leaving the dura intact. The implant was aligned to the craniotomy and fixed to the skull using a bone screw (F000CE094, Morris Precision Screws and Parts) placed 2-3 mm posterior to lambda. The implant periphery was glued to the skull (Vetbond, 3M) and cemented in place with dental cement (S380 S&B Metabond, Parkell Inc.). Following the cure of the cement, a custom, head-fixing titanium frame was fastened to the implant using three screws (3/32” flat head 0-80). A second application of dental cement enclosed the fastening screws. After surgery the mice recovered to an ambulatory state on a heating pad and then were returned to a clean home cage. Mice were administered meloxicam (2 mg/kg, s.c.) for three days and allowed a minimum of seven days to recover before any experimental procedures were initiated.

### Behavioral setup

Mice were housed in a reversed light-dark (12h-12h) room with experiments performed during the dark period, which is the normal waking and high activity phase of the circadian cycle in mice. After recovery from surgery, polymer window-implanted mice were habituated to the behavioral setup in increasing time increments (5 min, 15 min, 40 min, 1 hr) before experiments began. For the behavioral setup, mice were head-fixed on a low-friction, horizontal disk treadmill that allowed for natural movements such as walking and grooming (Fig. 1A). Once habituated, animals alternated between periods of awake quiescence (i.e., rest) and spontaneous walking, which were used for spontaneous locomotion analysis. To offset any potential effects created by the mildly curved path of the disk treadmill, a subset (22 of 62) of experimental sessions were recorded with the mouse head-fixed to the disk pointing in the opposite direction.

**Figure 1.**
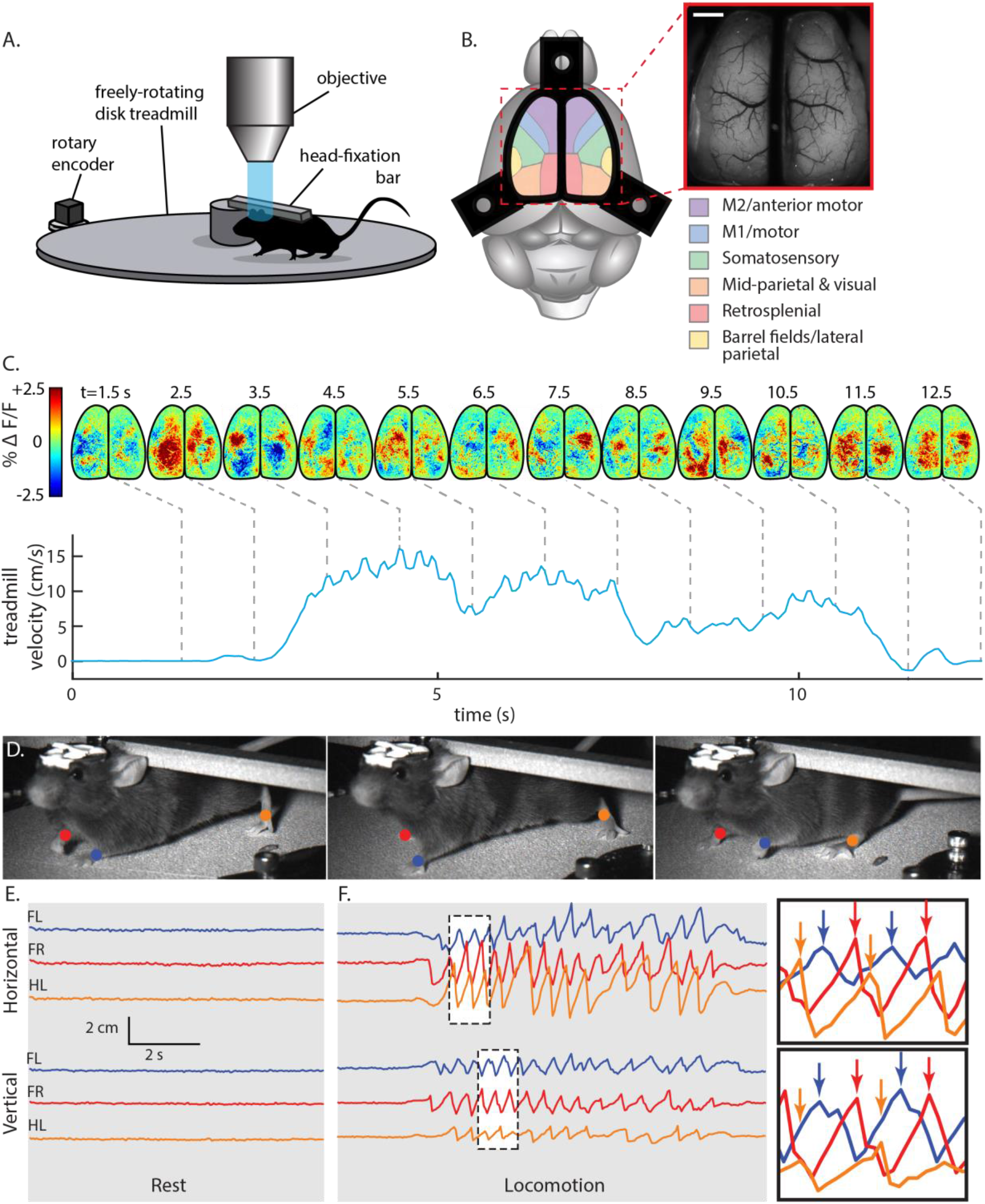
Mesoscopic cortical recording set-up and activity during locomotion. **A.** Experimental set-up for behavioral and Ca^2+^ fluorescence imaging of head-fixed mice. **B.** Approximate cortical regions, as defined by the Allen Brain Atlas Common Coordinate Framework (Allen Institute for Brain Science 2015), observable through the polymer skull (inset) by epifluorescence microscopy. Scale bar = 1 mm. Common Framework colors correspond to cortical regions defined in key. **C.** Treadmill velocity (bottom, blue line) is plotted for an example of spontaneous locomotion in a single mouse. Pre-processed maps of cortical activity (top) shown as mean subtracted % change in fluorescence (ΔF/F). For visualization only, frames were spatially filtered with a 3×3 moving mean and averaged across 5 frames. Grey dashed lines link maps of cortical Ca^2+^ fluorescence with time points prior to, during, and after locomotion. **D.** Markers used to track paw positions during periods of rest and locomotion for this example data. Blue, front left paw (FL); red, front right paw (FR); orange, hind left paw (HL). **E.** Horizontal (top) and vertical (bottom) paw displacements vs. time during rest. **F.** Horizontal (top) and vertical (bottom) paw displacements vs. time during locomotion. Maximum displacement of each paw (inset, arrows) show stereotypic, repeating step cycles.

Locomotion kinematics were calculated from the treadmill angular displacement as measured by a high resolution rotary encoder and recorded by an Arduino Uno microcontroller (Arduino) at 1 kHz. Velocity was determined and smoothed using a sliding average (100 time points, 1-time point step size). Locomotion was defined as periods of movement in which the wheel reached a velocity of 0.25 cm/s or greater. Working back from 0.25 cm/s, movement onset was then defined as the time wheel velocity first exceeded 0 cm/s, and movement offset was defined as the time velocity returned to 0 cm/s. Periods in which velocity remained between −0.25 cm/s and 0.05 cm/s were labeled as rest, while all remaining periods were discarded as “fidgeting” or backwards walking.

### Fluorescence imaging

Head-fixed mice were placed on the treadmill beneath a Nikon AZ-100 microscope (Fig. 1A). Single-photon fluorescence imaging was performed using a high-speed, Electron Multiplying CCD (Andor, iXon3) controlled with MetaMorph (Molecular Devices Inc.). A filter set with 480/20 nm excitation, 505 nm dichroic, and 535/25 nm emission filters was used (Chroma). Using the variable magnification function, the field-of-view was optimized to image the exposed dorsal cortical surface (6.2 mm x 6.2 mm) with a spatial resolution of 256 x 256 pixels (pixel size of ~41 μm x 41 μm). Images were acquired at 20 Hz, 20 ms exposure, for 5 mins (6000 frames), and 12 imaging trials were obtained in a session (i.e., the number of trials obtained in 1 day). Time between imaging trials ranged between 1-5 mins.

### Fluorescence imaging analysis

The Ca^2+^ fluorescence data from each imaging session was spatially registered using affine transformations. Consistent points on the visible blood vessels in the brain were manually selected and aligned using the built-in MATLAB function, *fitgeotrans*, with the “affine” method selected. All sessions were registered to the same representative session for a mouse. To remove motion artifacts within trials, all frames were registered to sub-pixel precision using the *dftregistration* MATLAB function (Guizar 2021). For each pixel, the time course of the fluorescence activity was then lowpass filtered with a 5^th^ order Butterworth filter with a cut-off frequency of 7 Hz. To remove artifacts potentially introduced through increased overall fluorescence or through blood flow and blood vessel constriction or dilation, masks were drawn over representative sections of background and blood vessels. The mean activity was taken from each of these masks, and the activity of each pixel was regressed against these traces. Only the residuals from these regressions were kept for further analysis, thus removing from the fluorescence signals contributions of background fluorescence and blood flow.

To reduce the dimensions of the data to a manageable level and decrease noise, we performed spatial independent component analysis (sICA) to identify a catalog of functionally relevant cortical regions. For each mouse, images from all trials were concatenated and compressed using singular value decomposition (SVD). Only the first 200 singular values were used to recreate the spatial dimension of the data (Musall et al. 2019). We computed the first 50 spatial independent components (ICs) using the Joint Approximation Diagonalization of Eigenmatrices (JADE) algorithm that decomposes mixed signals into ICs by minimizing the mutual information with a series of Givens rotations (Cardoso 1999; Makino et al. 2017; Sahonero-Alvarez and Calderon 2017). This method provides a blind segmentation of the cerebral cortex based only on statistical properties of the Ca^2+^ activity and does not use any prior assumptions regarding cerebral function or architecture. Masks of ICs were made by setting intensity values below 3.5 to 0. Masks covering less than 150 pixels or that, upon visual inspection, corresponded to artifacts not associated with cortical activity were discarded. This included vascular artifacts that survived the regression step above. An IC that included multiple discontinuous areas, such as homotopic cortical regions, was separated into individual ICs, and these individual ICs were used in subsequent analyses. All remaining IC masks were visually inspected, and any remaining area corresponding to blood vessels that were not separated from cortical activity were manually identified and removed. To group data across animals, ICs in each mouse catalog were manually assigned to 1 of 28 nodes of interest (14 per hemisphere) that were present in the majority of mice and corresponded, approximately, to known cortical regions based on the Common Coordinate Framework (Fig. 2)(Allen Institute for Brain Science 2015).

**Figure 2.**
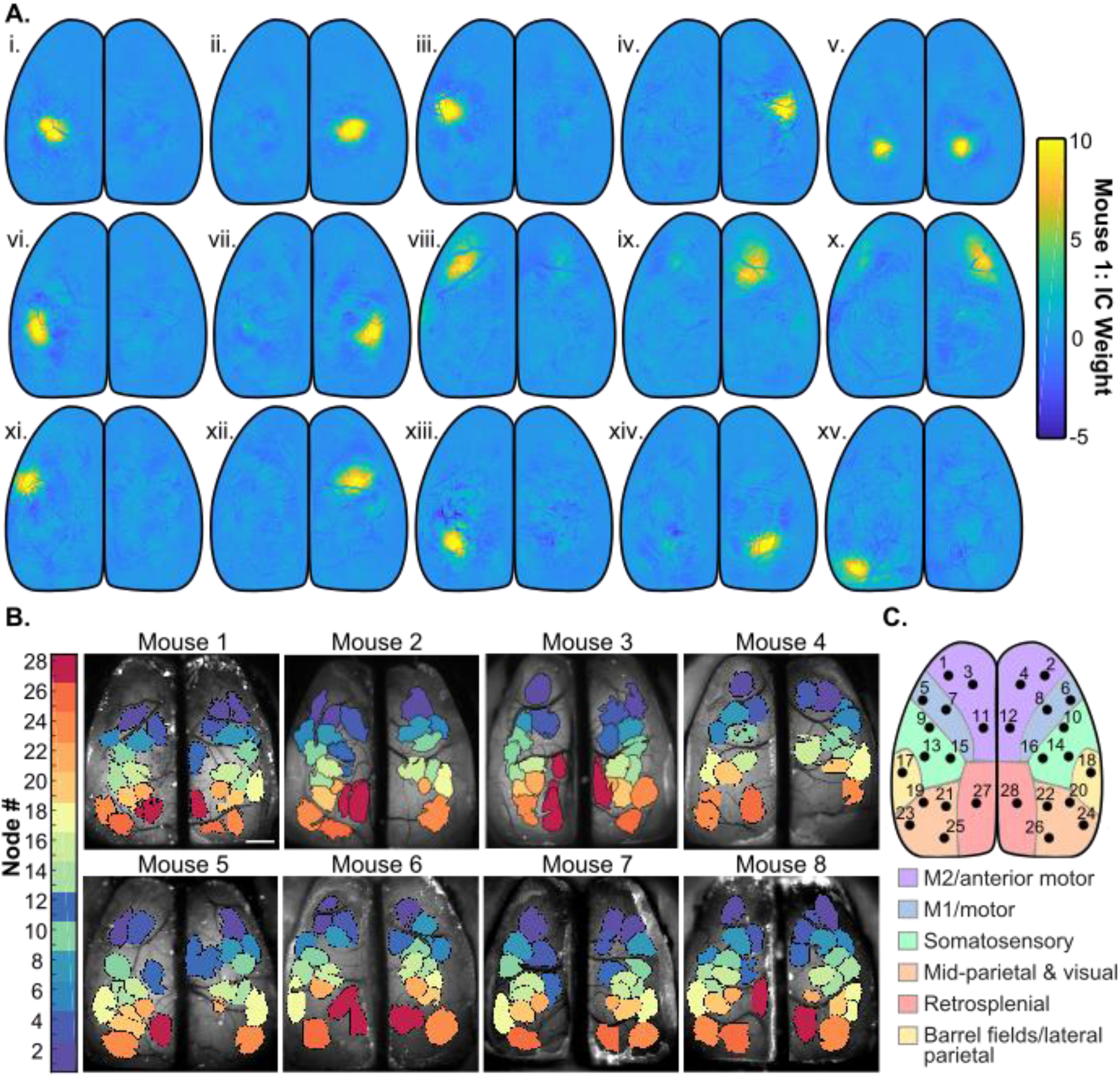
Spatial segmentation of the dorsal cerebral cortex using sICA. **A**. 15 ICs representing cortical activity from mouse 1 calculated using JADE ICA, before thresholding and manual artifact removal. **B**. IC catalogs of each mouse after thresholding and manual artifact removal. Color corresponds to the node identity assigned to each one of the 28 common ICs found across mice as shown in C. **C.** Locations of the 28 common ICs that define the network nodes observed across mice mapped onto the Common Coordinate Framework (Allen Institute for Brain Science 2015).

### Behavior Periods

Recordings were divided into 6 behavior periods, each 3sin duration, as defined by treadmill velocity (see Fig. 3B and 4A): 1) rest (see definition above); 2) pre-locomotion (rest just prior to locomotion onset; “prep.”); 3) initiation of locomotion (locomotion just after locomotion onset; “init.”); 4) continued locomotion (periods of steady-state, continued locomotion outside of transition periods; “cont.”); 5) termination of locomotion (locomotion just prior to locomotion offset; “term.”); and 6) post-locomotion (rest just after locomotion offset; “after”). Periods of rest or continued locomotion that lasted longer than 3s were divided into multiple 3s segments, and remainder data at the end of the period was removed. Periods less than 3s were also removed. We chose 3s periods (60 time points) because this provides sufficient data to calculate robust Pearson correlations on the associated fluorescence time series for the functional connectivity analysis (see below).

**Figure 3.**
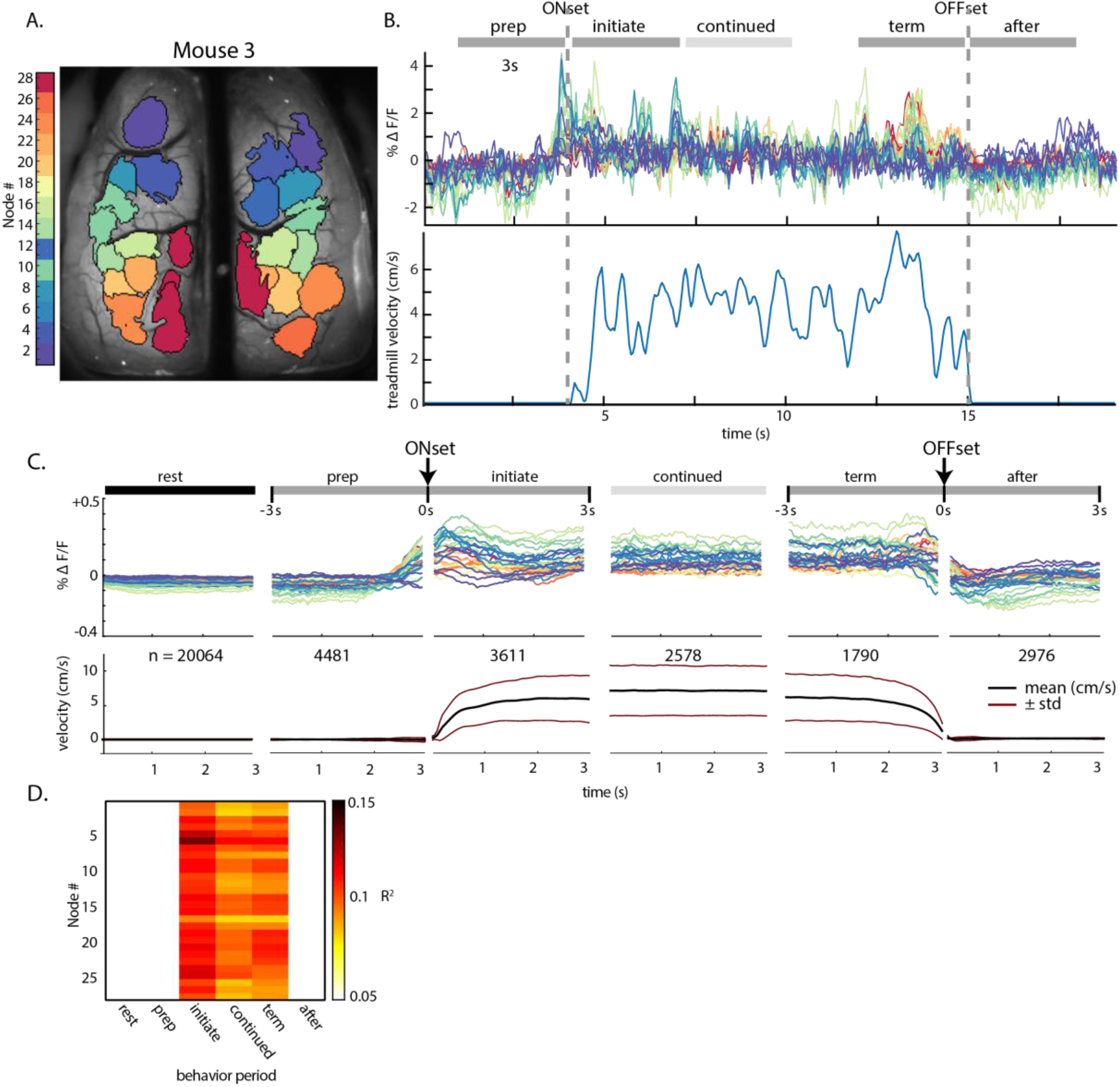
Fluorescence activity (ΔF/F) within ICs across behavior periods. **A.** IC catalog of mouse 3, as shown in Figure 2B. **B.** Top: Example fluorescence time series from each IC for a single bout of locomotion in mouse 3, with the timeline of the 6 behavior periods on top. Bottom: corresponding treadmill activity. **C**. Top: Fluorescence activity from the common set of 28 nodes averaged over all mice and across all instances of each 3 s behavior period (n = number of periods averaged). Abbreviations: pre., pre-locomotion; initiate., initiation of locomotion; continued, continued locomotion; term., termination of locomotion; after, after locomotion. Bottom: Mean (black lines) and s.d. (red lines) of treadmill velocity across each behavioral time period for all mice. **D.** Average R^2^ values from regressions of fluorescence activity against treadmill acceleration and velocity for each behavior period. Only significant R^2^ values are shown.

**Figure 4.**
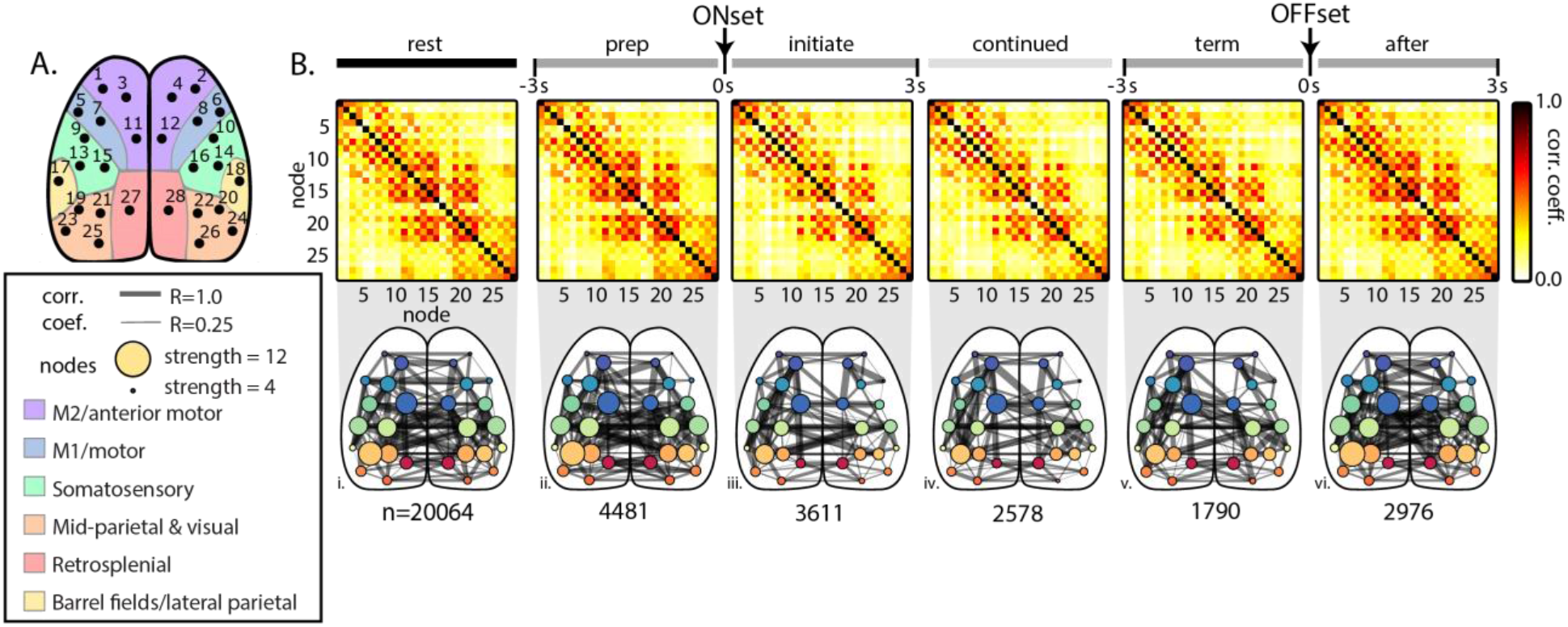
Functional connectivity among the common cortical nodes during behavior periods across all mice. **A.** Diagram of nodes common to the catalog ICs across mice mapped onto the Common Coordinate Framework (Allen Institute for Brain Science 2015). Region color corresponds to key in Fig. 2B. **B** Top: Averaged correlation matrices between nodes across all mice. Bottom: Graphical representation of the correlation matrices shown above superimposed on the dorsal cortical surface. Width of connecting edges indicates the Pearson correlation coefficient (ρ ≥0.25). The size of the node reflects the strength; that is, the sum of all correlations with that node. n=number of 3s time periods included in each behavior period across animals.

### Regression of fluorescence activity against treadmill speed

The masks from the IC catalog for each mouse were used to extract mean fluorescence time series (based on the preprocessed series of image data), and the resulting time series were linearly detrended. To determine how fluorescence activity in each node modulated with the parameters of locomotion, a multiple linear model was used to regress the fluorescence activity of each node to treadmill velocity and acceleration, using the MATLAB built-in *regress* function. R^2^ values were averaged for each behavior period over all trials and animals.

### Functional connectivity analysis

For each 3s period, the correlation coefficients (ρ) were calculated between the time series from all catalog ICs to generate a correlation (i.e., adjacency) matrix, and these matrices were averaged to create mean adjacency matrices for each behavior period. To combine data across animals, the averaged correlations were assigned to the appropriate corresponding node. If more than one catalog IC was assigned to a node, then the correlation values of those catalogue ICs were averaged. The mean adjacency matrices were then averaged across the subjects. For each pair of regions, the significant change in correlation of activity between behavior periods was calculated using custom MATLAB code. The difference in correlation was compared to a null distribution of differences from 500 reshufflings of the correlations across behavior. Significance was determined by α<0.05 and corrected with the false discovery rate (Genovese et al. 2002).

To further quantify the functional relationships between brain regions during locomotion, the network centrality of ICs was calculated on 3s correlation matrices using MATLAB code from the Brain Connectivity Toolbox (Rubinov and Sporns 2010). The eigenvector centrality was calculated from the correlation adjacency matrices for each 3s period and averaged within each mouse before being averaged across mice. Similar to the comparisons among the correlations between regions, the difference in eigenvector centrality for each region from one period to another was compared to a null distribution of differences from 500 reshufflings of the centralities across behavior. Again, significance was determined by α<0.05 and corrected with the false discovery rate.

Since it is possible that large, global increases or decreases in fluorescence activity could increase correlation coefficients between nodes and obscure more subtle changes in connectivity, an additional analysis was performed in which low frequency changes in fluorescence activity were removed using a 5^th^ order high pass Butterworth filter with a cutoff filter of 2 Hz. The filter was applied to the mean fluorescence time series extracted from the IC masks for each mouse. Correlation and eigenvector centrality calculations were repeated on the filtered data as described above.

### Granger causality analysis

Granger causality among the nodes was determined as a measure of the directional influence between cortical regions during different behavioral periods. We used the Multivariate Granger Causality MATLAB Toolbox (Barnett and Seth 2014), with the ordinary least squares model estimation and regression information criteria and the Akaike information criterion (AIC) model order. The model was limited to a maximum of 20 lags. For each animal, the time series from all instances of a 3s behavior period were inputted to the algorithm as separate trials to generate a single, multivariate causality adjacency matrix for each mouse. These individual adjacency matrices were averaged across mice as described above. Granger causality estimates the causality between time series in both directions. In order to have a single directional value representing the net relationship between two regions, a “total causality” value was calculated based on the difference in magnitude of the corresponding causalities. In adjacency matrices, total causality direction is indicated as a positive or negative value. Significant changes in causality between behavioral periods were determined similar to the approach used for significant changes in correlation and centrality. The difference in causality and total causality from one period to another between each pair of ICs was compared to a null distribution of differences from 500 reshufflings of the causalities across behavior. Significance was determined by α<0.05 and corrected with the false discovery rate.

### Hemodynamic correction

Blood flow increases with neuronal activation, and oxygenated blood absorbs light with peak absorption at ~530 nm, decreasing the duration of the increased GCaMP fluorescence (Ma et al. 2016). Therefore, additional experiments were performed to evaluate the effects of hemodynamics and other Ca^2+^ -independent fluorescence changes such as flavoprotein autofluorescence (Vanni and Murphy 2014; Jacobs et al. 2018), using 3 mice (1 from the original cohort and 2 additional). Data was collected in 5-minute stacks, similar to the primary dataset. Data was collected in 36 stacks across 3 recording days for mouse #8, 35 stacks across 8 days for mouse #9, and 68 stacks across 12 days for mouse #10, for a total of 11.58 hours of data. We used dual-wavelength illumination to capture both Ca^2+^ -dependent (470 nm, blue light) and Ca^2+^- independent (405 nm, violet light) GCaMP6f signals on consecutive frames using a Cairn OptoLED driver (Cairn OptoLED, P1110/002/000; P1105/405/LED, P1105/470/LED) (Ma et al. 2016; Allen et al. 2017; Jacobs et al. 2018; Musall et al. 2019; MacDowell and Buschman 2020). An excitation filter (ET480/40, Chroma) was placed in front of the 470 nm LED, then both light sources were combined into the parallel light path of a Nikon AZ100M macroscope through a dichroic mirror (425 nm, Chroma T425lpxr), which was reflected off a second dichroic (505 nm, Chroma T505pxl) to the brain. Cortical GCaMP6f emissions then passed back through a second dichroic into a sCMOS camera (Andor Zyla 4.2 Oxford Instruments). Exposure times for each frame was 18ms, synced via TTL pulses from a Cambridge Electronics 1401 (Cambridge Electronic Design Limited) acquisition system that controlled both LEDs and the external trigger of the Andor Zyla 4.2. Frames were captured at 40 Hz (20 Hz per channel) at 256 x 256 pixels per image.

Using a previously described correction method (Ma et al. 2016; Jacobs et al. 2018; MacDowell and Buschman 2020), the Ca^2+^-independent signals were removed by first calculating the per-pixel average intensity in both channels, then scaling the 405 nm channel to a similar level of the 470 nm channel by multiplying by the ratio of the per-pixel averages. The scaled 405 nm signal was then subtracted from the 470 nm signal and the resulting signal was then normalized by dividing by the scaled 405 nm signal. All subsequent processing, including sICA, functional connectivity and network measures, was identical to that preformed on the mono-wavelength, uncorrected data.

All MATLAB analysis code including for sICA segmentation of mesoscale Ca^2+^ imaging; eigenvector centrality; and hemodynamic correction are available upon request.

## RESULTS

### Database

We collected imaging data from eight GCaMP6f transgenic mice. Eight imaging sessions were performed on seven of the animals, while six imaging sessions were performed on the remaining animal, resulting in a total of 62 sessions. During each imaging session, we obtained 60 min of data (12 trials x 5 min each), resulting in 480 min of imaging in 7 animals and 360 min in 1 mouse. Mice spent an average of 75.7 ± 10.5% of recording time at rest, 21.3 ± 10.4% locomoting, 2.2 ± 0.9% fidgeting, and 0.9 ± 0.4% of recording time moving backwards. Across the 8 mice, we fully analyzed a total of 2,743 min of awake quiescence and 776 min of spontaneous locomotion.

### Changes in Ca^2+^ fluorescence during behavior

Mice were head-fixed over a disk treadmill that allowed for spontaneous locomotion (Fig. 1A), and wide-field Ca^2+^ imaging performed through an implanted polymer skull (Ghanbari et al. 2019) (Fig. 1B). Regions across the dorsal cerebral cortex show dynamic changes activity as the mouse transitions from rest to locomotion and back to rest (Fig. 1C). For each animal, spontaneous locomotion is characterized by an alternating and rhythmic step cycle typical of coordinated walking (Fig. 1D-F).

### Functional segmentation of the cortex

We segmented cortical activity into functionally distinct regions based on changes in Ca^2+^ fluorescence using sICA, a blind source separation tool commonly used in fMRI studies (Cardoso 1999; Makino et al. 2017; Sahonero-Alvarez and Calderon 2017). By minimizing the mutual information between regions to segment the cortex, sICA does not make assumptions about the underlying neuroanatomy of individual animals. The processed fluorescence datasets yield 22-31 spatial ICs per mouse, with 28 ICs in common across mice. As an example, 15 individual ICs from a single mouse are shown in Fig. 2A. The entire, thresholded IC catalogs for each animal are shown in Fig. 2C. Across the IC catalogs, 28 ICs were the most consistent and approximately equivalent across the majority of the 8 mice (Fig. 2B and C). These 28 common ICs were mapped onto the Common Coordinate Framework (Fig. 2B) and are the nodes in the following cortical network analyses. Though there are considerable similarities in the IC catalogs across the 8 mice, individual differences in functional anatomy are preserved with this method.

### Fluorescence activity during behavioral transitions and regression analysis

To analyze the functional connectivity of the cortex during the transitions between rest and locomotion, we extracted the average of the fluorescence time series from each IC in relation to the 6 behavioral periods (see Methods), as shown for an individual bout of walking in a mouse (Fig. 3A) and for the average data across all animals (Fig. 3B). At the transition from rest to walking, most nodes exhibit an increase in activity prior to locomotion that peaks around locomotion onset (Fig. 3C). The increases in fluorescence include nodes in primary motor regions, primary sensory areas (including somatosensory and visual cortices) and throughout posterior parietal and retrosplenial areas. This agrees with previous reports of motor, somatosensory, parietal, auditory, retrosplenial, and visual cortical engagement during locomotion (Drew et al. 2008b; Niell and Stryker 2010; Petersen et al. 2012; Saleem et al. 2013; Favorov et al. 2015; Drew and Marigold 2015; Schneider and Mooney 2018; Clancy et al. 2019) and highlights the involvement of multiple cerebral cortical regions in processing and integrating information related to locomotion. In all regions except the most anterior nodes, this increase begins 784 ± 201 ms before the onset of locomotion by an increase greater than the mean + 2.5 s.d of the fluorescence at rest. Nodes 1 and 2, in contrast, decrease in mean fluorescence. Throughout locomotion, average neural activity remains elevated compared to rest and, and decreases to baseline levels on return to rest (Fig. 3C). This pattern of fluorescence changes was present in all mice.

To quantify the degree to which the parameters of locomotion are represented in the Ca^2+^ signals, the fluorescence activity of each node was regressed to treadmill velocity and acceleration, revealing widespread encoding of these parameters across the cerebral cortex (Fig. 3D). The strongest correlations occur during the initiation period, with somewhat smaller R^2^ values during continued and termination periods. During these three periods, the R^2^ values for all nodes reached significance (F(2, 58) >3.16, α <0.05). Conversely, the regressions were not significant for the other periods. Importantly, the encoding occurs across all nodes, consistent with a global representation of parameters of locomotion throughout the dorsal cerebral cortex.

### Correlations between nodes

To understand the functional coupling among the ICs during these transitions, we defined six behavioral periods related to locomotion, each 3s in length: 1) rest, 2) pre-locomotion, 3) initiation of locomotion, 4) continued locomotion, 5) locomotion prior to termination, and 6) rest after termination of locomotion (see Methods and Fig. 3A and C). Within each 3s behavioral period for a mouse, the Pearson correlation (ρ) was calculated between the fluorescence time series for each possible pair of ICs. The resulting correlation coefficients were averaged across all instances of the behavior period in that mouse and then averaged across mice using the 28 common nodes. Each node’s mapping to the Common Coordinate Framework is shown again in Fig. 4A. The average correlation coefficients between nodes were plotted as both correlation matrices and network graphs (Fig. 4B). Across behavior periods, nodes that are anatomically adjacent tend to show high correlation. For example, nodes located in the parietal cortex (9-22) exhibit especially high connectivity, except for nodes 17 and 18 in the lateral parietal area that are approximately equivalent to the barrel fields. Similarly, the nodes in the premotor, primary and visual cortices have high internal functional connectivity within and across hemispheres. Qualitatively, there is a general decrease in functional connectivity during movement initiation that persists through the termination of locomotion, followed by an increase in correlations after movement.

We used a permutation test with false discovery rate correction to evaluate the significant changes in mean correlations between nodes, both increases and decreases, between behavior periods (Fig. 5). Compared to rest, continued locomotion exhibits both significant decreases and increases in correlation depending on the nodes and cortical regions (Fig. 5A and B). Unexpectedly, nodes within M1 (5-10) decrease in correlation with most other regions including with homologous nodes in the contralateral hemisphere (82 of a possible 162 connections). However, there are some increases in correlation among nodes within M1 (12 of a possible 30 connections). Also, nodes within the primary somatosensory cortex exhibit decreases in correlations widely. There are two prominent patterns of increased functional connectivity. First, nodes (1-4) in M2 show increases in connectivity with visual and retrosplenial regions as well as increases with other parietal regions (49 of 108 possible connections). Second, retrosplenial nodes 27 and 28 exhibit large increases with regions across the cortex (35 of 54 possible connections).

**Figure 5.**
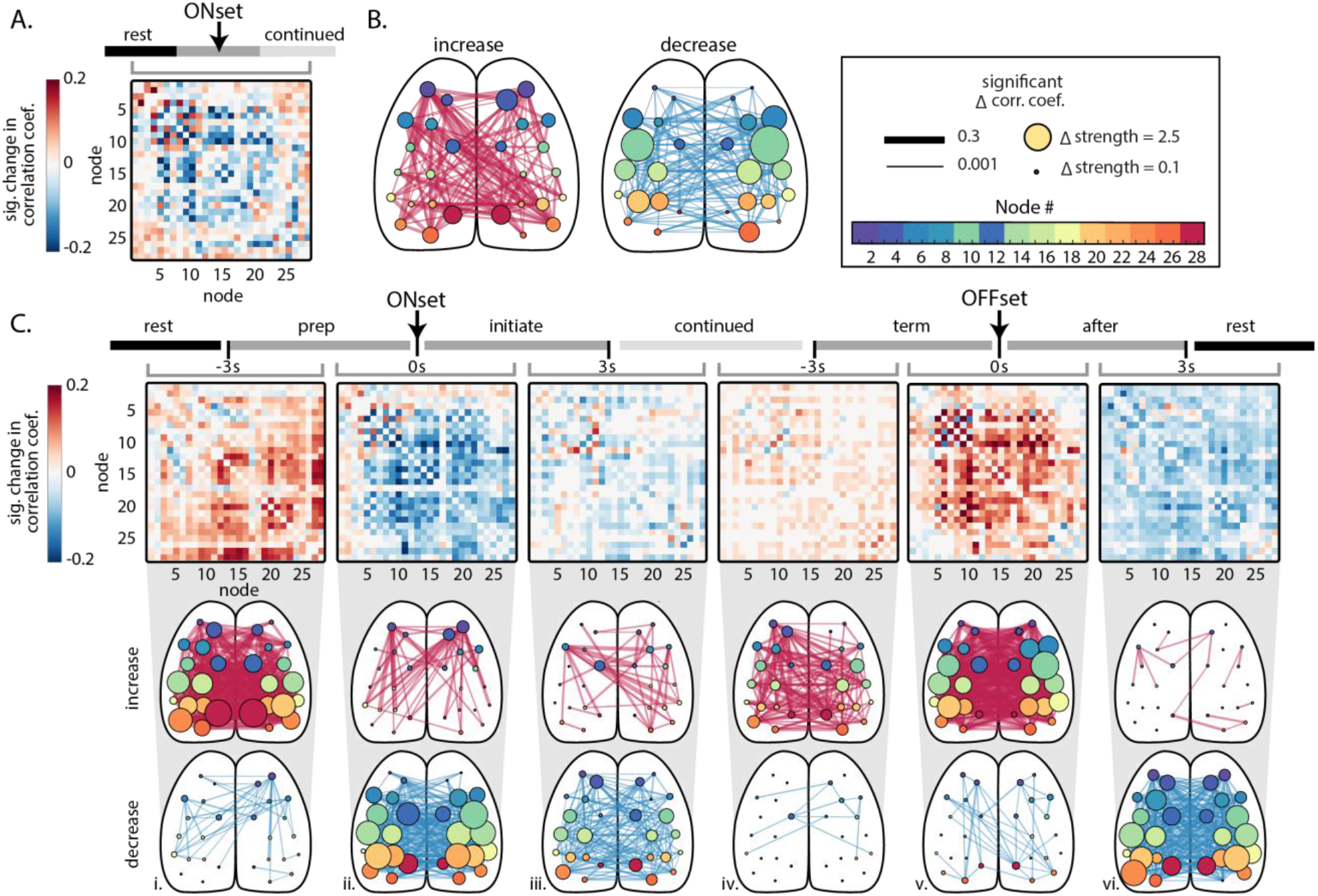
Significant changes in correlations between nodes across behavioral periods. **A.** (left) Matrix of significant changes in correlation between nodes comparing rest to continued locomotion (α<0.05, permutation test with false discovery rate correction). **B.** Significant increases (center) and decreases (right) from rest to continued locomotion shown in graph representations, superimposed on the cortical surface. The size of each node reflects the magnitude of significant increase or decreases, respectively, for that region. **C.** Significant changes in correlations across sequential behavior periods, calculated and displayed as in A and B. Directly above each correlation matrix, the grey bracket indicates the adjacent behavior periods being compared (i-vi).

Next, we evaluated the significant changes in correlations across sequential behavior periods. From rest to the pre-locomotion period, functional connectivity increases between many nodes, with large increases in the nodes located in visual and somatosensory regions to those in other regions, and the strongest increases involve the retrosplenial nodes (Fig. 5Ci). In contrast, correlations between the anterior M2 nodes and many other nodes significantly decrease (18 of 54 possible connections). Nodes within M1 (5 and 6) show decreases in correlation with somatosensory nodes (9, 10 and 13-16; 8 of 12 possible connections). During initiation of locomotion compared to pre-locomotion, correlations increase between the two most anterior nodes (1 and 2) in the M2 region and nodes in nearly all other regions (Fig. 5Cii; 32 of 54 possible connections). However, the correlations between anterior M2 nodes (1 and 2) and primary motor nodes (5-10) significantly decrease (8 of 12 possible connections). Correlations decrease between nodes in most other regions, with primary motor, somatosensory and parietal nodes exhibiting large decreases. As the animal transitions from the initiation of locomotion to continued locomotion, correlations across much of the dorsal cortex continue to decrease (Fig. 5Ciii). These patterns reverse as the animal progresses through the termination of locomotion and post-locomotion periods to rest (Fig. 5Civ-vi), with striking increases in functional connectivity across the dorsal cortex with the termination of locomotion and a widespread decrease as the animal returns to rest.

It is possible that the large shift in overall fluorescence activity occurring during different behavioral periods (Fig. 3C, top) may be the main contributor to the changes in correlations between nodes and may obscure more subtle changes in connectivity. To investigate this possibility, we removed these low frequency fluorescence shifts by applying a high pass filter with a 2 Hz cutoff frequency to the average IC time series extracted from the fluorescence images (Supplemental Fig. S1). Filtering eliminates the large magnitude changes in fluorescence at the start and the termination of locomotion (Supplemental Fig. S1A). Notably, the patterns of changes in correlations across behavior periods remain intact, as observed in the adjacency matrices and network diagrams (Supplemental Fig. S1B-E), both for the comparison of rest versus continued locomotion (Supplemental Fig. S1B and D) and for comparisons between adjacent behavior periods (Supplemental Fig. S1C and E). Therefore, global changes in fluorescence do not dictate the dynamic patterns of functional connectivity that occur from rest to locomotion and with return to rest.

### Eigenvector centrality during behavioral transitions

To further characterize the changes in network structure, we calculated the mean eigenvector centrality of nodes within each behavior period and compared how centrality changed across periods. Eigenvector centrality provides a measure of how tightly connected the behavior of a node is to all other nodes in the network (Rubinov and Sporns 2010). The average eigenvector centrality across all mice, for each node and behavioral period (Fig. 6A), shows the overall importance of different nodes. For example, somatosensory and parietal nodes show high centrality during rest, preparation and after periods. During locomotion periods, the centrality of these regions generally decreases with other nodes increasing in importance. Therefore, we determined the significant changes in centrality from rest to continued locomotion and between adjacent period by calculating a permutation distribution with false discovery rate correction.

**Figure 6.**
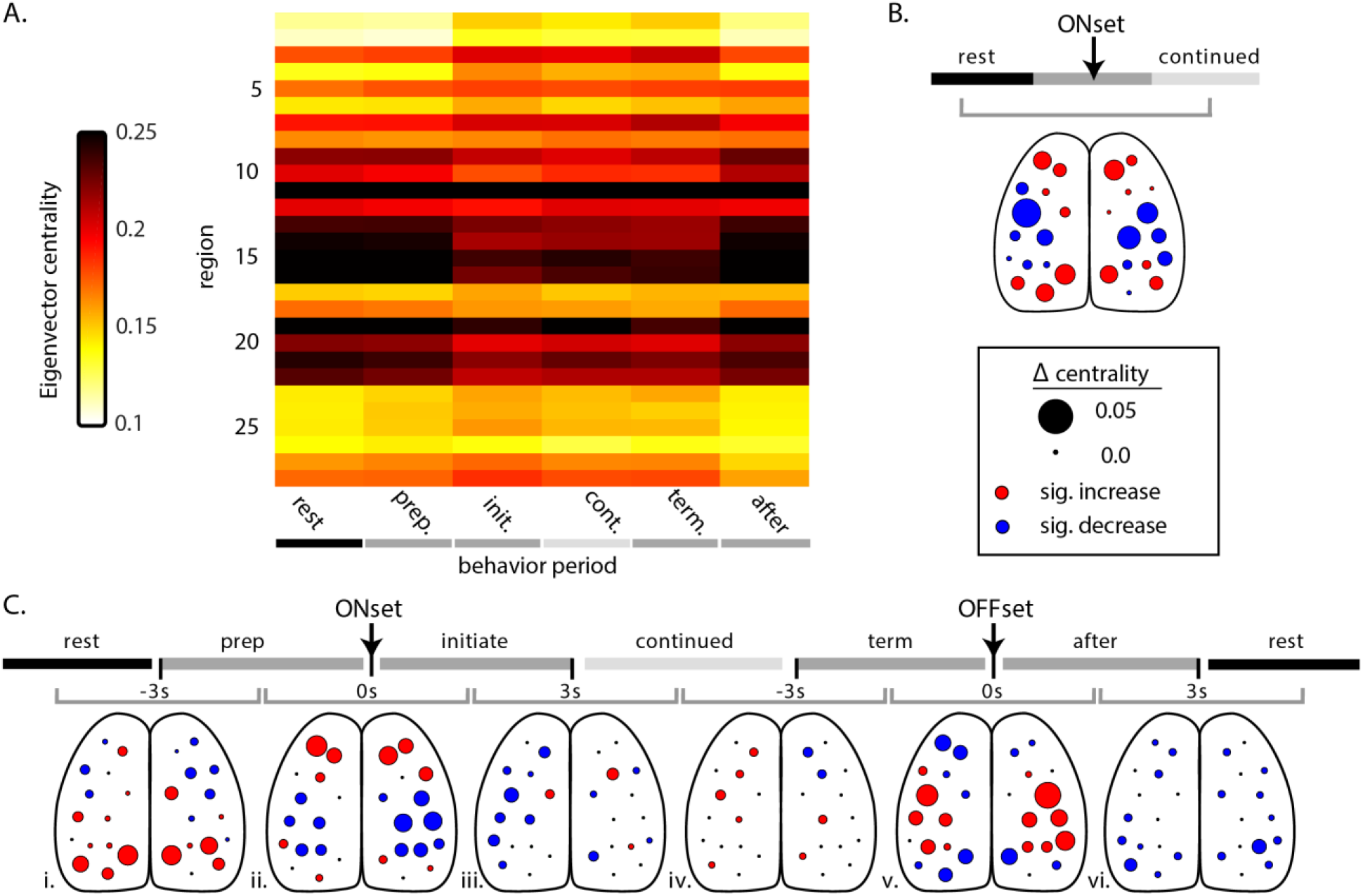
Eigenvector centrality across behavioral periods. **A.** Average eigenvector centrality for each node during the behavior periods, across mice. **B.** Significant change in centrality comparing rest to continued locomotion across all mice. Size of circles indicate magnitude of the change, while circle color depicts the direction of significant change (red=increase, blue=decrease, black=not significant; α<0.05, permutation distribution with false discovery rate). **C.** Significant change in centrality across sequential behavior periods (shaded timeline on top). As in Figure 5, grey bracket indicates the adjacent behavior periods being compared (i-vi). Color and size of circles as described in B.

Comparing rest with continued locomotion (Fig. 6B) reveals robust changes in centrality that confirm the changes observed in the correlations (Fig. 5B). In M2, nodes 1-4, 11, and 12 increase in eigenvector centrality, with the largest in nodes 1-4. Nodes 5-8 in the M1 exhibit a mixture of smaller increases and larger decreases. Centrality decreases in somatosensory and more posterior parietal nodes (9-22), with the largest decreases occurring in somatosensory cortex nodes (9, 15, 10 and 16). Nodes 23-25 in the visual areas and 27-28 in retrosplenial areas increase in centrality when comparing rest to continued locomotion.

Comparing adjacent behavior periods, there are significant increases in centrality of visual and retrosplenial nodes as the animal transitions from rest to pre-locomotion (Fig. 6Ci). Meanwhile, increases in premotor nodes (1-4) and decreases in somatosensory and middle parietal nodes (9-22) do not occur until the initiation of locomotion (Fig. 6Cii). The centrality changes reverse during the transition from the termination of locomotion to post-locomotion with large increases in nodes in the somatosensory and parietal areas and decreases in premotor, retrosplenial and visual nodes (Fig. 6Cv.). This is followed by more general decreases in centrality as the animal returns to rest (Fig. 6Cvi). As observed for significant changes in correlation, the changes in eigenvector centrality are also preserved following high-pass filtering (Supplemental Fig. S1F and G) and demonstrate centrality is independent of the large fluorescence shifts that occur during locomotion. These changes in network centrality highlight the dynamic roles played by different functional regions and their interactions between behavior periods.

### Granger Causality

The Granger causalities between all nodes were calculated to estimate the directionality of the functional connectivity during different behavior periods (Fig. 7A). Granger causality was calculated for each direction of every possible pair of nodes (i.e. time series X Granger causing time series Y as well as time series Y Granger causing time series X), plotting the resultant causality in both directions (Fig. 7A). There is a tendency for certain nodes, for example 1 and 4 in the premotor region and 26 in the visual region, to causally impact several other nodes in the cortex. Conversely, some nodes receive input from many others. Qualitatively, causality increases from rest to pre-locomotion, continues to increase during continued locomotion and increases even further at the termination of locomotion (Fig. 7A).

**Figure 7.**
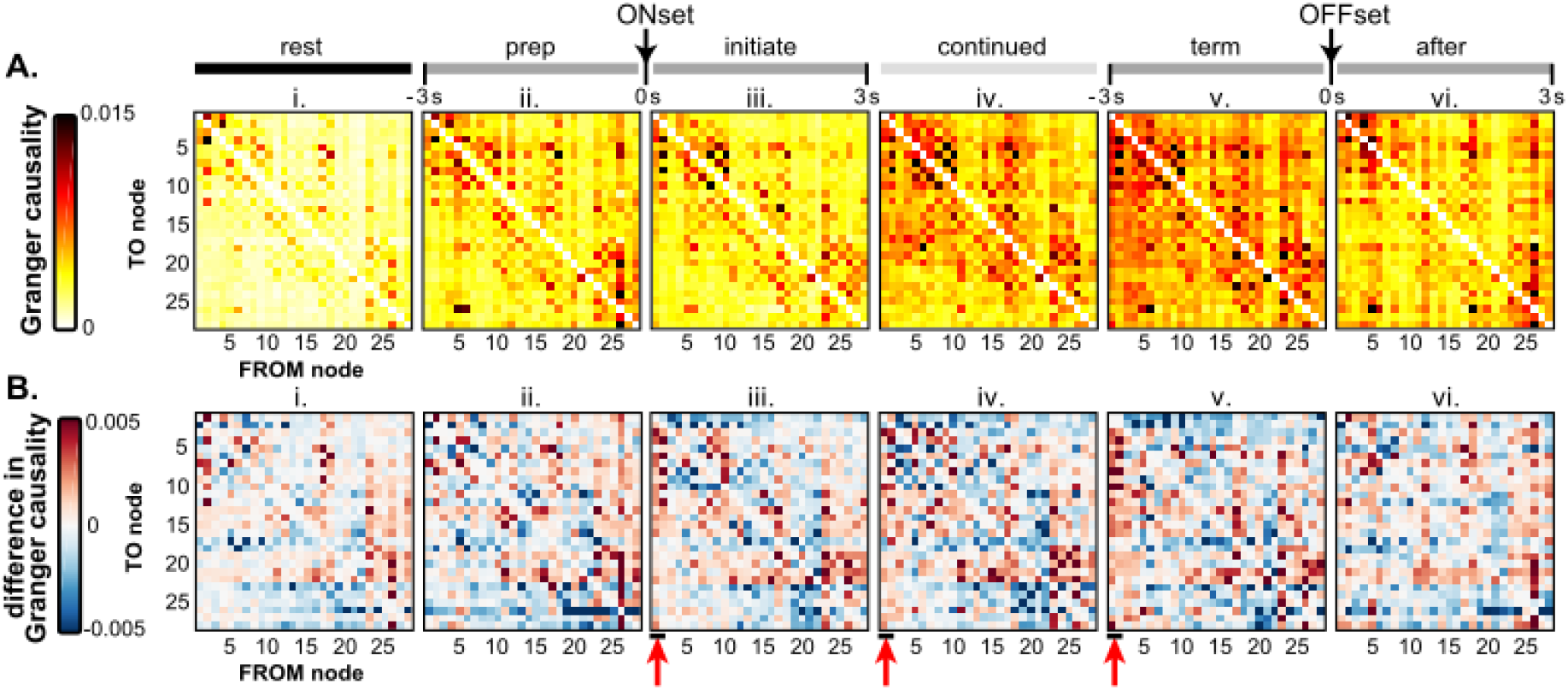
Granger causality between nodes during the behavior periods. In these plots, rows represent the TO node Granger causality, and columns represent the FROM node causality. **A.** Average Grainger causality between nodes in both directions for each period, across all mice. **B.** Total Granger causality between nodes determined by subtracting the upper triangle of the matrix from the lower triangle. Red Arrows highlight the outward causality of anterior M2 nodes (nodes 1-2) during locomotion.

To simplify, we determined a single total causality vector between pairs by summing the causalities between each pair of nodes. The magnitude and direction (positive or negative) of the total Granger causality “to” and “from” each node is shown in Fig. 7B. During all behavior periods, visual nodes (23-26) exert greater Granger causality (positive values) on other nodes than others exert back on visual nodes (Fig. 7Bi-vi). A similar pattern is observed for nodes 17 and 18, which correspond approximately to the barrel fields, suggesting these two sensory systems exert net positive Granger causality on many other cortical areas. Most notable, however, nodes 1 and 2 in the anterior M2 area exert Granger causality on many other nodes beginning during the initiation of locomotion (Fig. 7Biii, 44 of 54 possible connections) and persisting through continued (44 of 54 possible connections), and termination (47 of 54 possible connections) (Fig. 7Biv, v), suggesting the premotor cortices play important causal roles from the start to end to locomotion (see red arrows in Fig 7Biii-v).

We calculated the significant change in Granger causality between adjacent behavior periods for both directed (Fig. 8A) and total causality (Fig. 8B). Comparing locomotion to rest, the changes include scattered increased influence among nodes located in premotor, primary motor and somatosensory regions (Fig. 8A and C). Most of the significant changes between behavior periods occur across locomotion onset and across locomotion offset. The most prominent changes in Granger causality are increases from the anterior M2 nodes, from preparation to initiation of locomotion (Fig. 8C and D; 10 of 54 possible connections in C, 19 of 54 in D; see red arrows). This finding supports the average causalities (Fig. 7A) as well as the increases in correlation (Fig. 5Cii) and centrality (Fig. 6Cii) and suggests that anterior M2 nodes begin exerting a causal influence on other cerebral cortical regions at the start of locomotion and continue to do so until locomotion ends.

**Figure 8.**
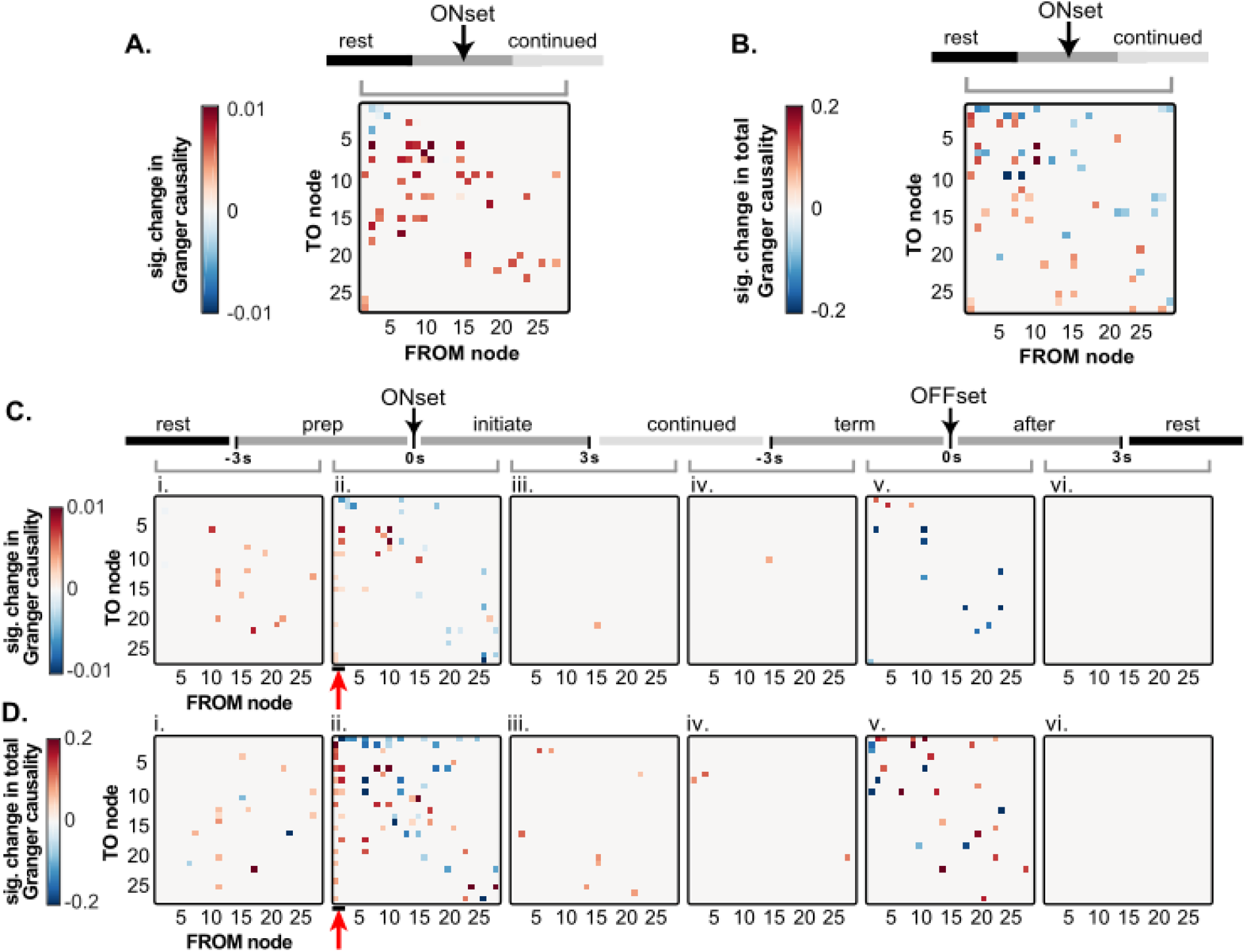
Significant changes in Granger causality between nodes across behavioral periods In these plots, rows represent the TO node Granger causality, and columns represent the FROM node causality. **A.** Significant change in Granger causality in both possible directions from rest to continued locomotion. **B.** Significant change in total Granger causality, as shown in Fig. 7B, from rest to continued locomotion. **C.** Significant change in Granger causality across sequential behavior periods **D.** Significant change in total Granger causality across sequential behavior periods. (Significance based on α<0.05 using permutation distribution with false discovery rate). Red arrows highlight the significant increase in outward causality of anterior M2 nodes (nodes 1-2) at the onset of locomotion. Grey bracket indicates the adjacent behavior periods being compared (i-vi) in C and D.

### Hemodynamic contributions

We imaged 3 mice using dual-wavelength imaging to correct for Ca^2+^-independent changes in fluorescence to remove the hemodynamic component from the GCaMP signal (see Methods). The IC catalogs were similar to those in the uncorrected dataset and were grouped into the same 28 common nodes (Supplemental Fig. S2A and B). The overall pattern of changes in fluorescence across the behavior periods (Supplemental Fig. S2C) were similar to those observed in the original 8 mice (see Fig. 3C). Because less data was used, these catalogs included fewer ICs than were found in the primary dataset, although all but node 11 were represented in the 3 mice at least once. As in the primary dataset, anterior M2 nodes increased in correlation with the majority of more posterior nodes comparing rest to continued locomotion (Supplemental Fig. S2D and F, left). These increases occurred at the onset of locomotion, from the preparation to initiation periods (Supplemental Fig. S2Eii and Gii), and then the correlations decreased to rest levels at the offset of locomotion, from termination to after periods (Supplemental Fig. S2Ev). These changes in correlation were confirmed in the changes in centrality (Supplemental fig. S2G). These data demonstrate that the blood flow had minimal effects on the sICA segmentation and network connectivity during the transitions between rest and locomotion.

## DISCUSSION

We performed wide-field Ca^2+^ imaging in mouse cerebral cortex during locomotion and studied the functional connectivity between cortical regions. The results confirm previous studies demonstrating that locomotion activates many regions in the cerebral cortex (Niell and Stryker 2010; Ghose 2015; Dadarlat and Stryker 2017). Three key findings resulted from this study. First, correlations and centrality of nodes in the primary motor and somatosensory cortices decrease during locomotion compared to rest. Second, there is an increase in correlations and centrality of the retrosplenial cortex during locomotion. Third, the correlations, centrality, and the outward causality of nodes in the anterior M2 region increase at the onset of locomotion. Notably, the changes in network connectivity are preserved if the large shifts in Ca^2+^ fluorescence levels are removed, indicating these changes in the network are independent of global, low frequency shifts in cortical excitability.

### Cortical segmentation with sICA

In the present study, sICA uncovered 22-31 independent spatial regions per mouse that we used to functionally segment the cerebral cortex and characterize the changes in Ca^2+^ epifluorescence. A previous implementation of sICA using JADE (Makino et al. 2017) obtained 16 ICs of approximately similar size and locations across the dorsal cerebral cortex. Our greater number of ICs can likely be attributed to the higher sensitivity of imaging through transparent polymer windows as opposed to thinned bone, as well as differences in data collection, filtering, and analyses. The minimization of mutual information between regions by sICA provides a complementary approach to that provided by the Common Coordinate Framework (Allen Institute for Brain Science 2015). Here, we elected to determine the ICs independently and then align the ICs with the major cortical divisions in the mouse. Our implementation of sICA was performed over all available data from a given mouse, which allowed for determination of the ICs over a large data base (432,000-576,000 images). Other segmentation approaches used with wide-field Ca^2+^ imaging include seed-based correlation (Vanni et al. 2017), seed-correlation co-localized with tract-tracing (Mohajerani et al. 2013), spike-triggered mapping (Xiao et al. 2017), and non-negative matrix factorization (MacDowell and Buschman 2020). While there have been few direct comparisons of these methods in Ca^2+^ imaging, different analytical approaches will likely provide complementary information.

### Hemodynamic correction

While the analysis removed as much blood flow contribution to the signal as possible by regressing out fluorescence activity correlated with blood vessels, some fluorescence changes resulting from hemodynamic activity may have remained. In a group of 3 animals, we determined the influence of increased blood flow on the spatial ICs and the functional connectivity network, using dual wavelength imaging (470 and 405 nm) to remove Ca^2+^-independent changes(Vanni and Murphy 2014; Jacobs et al. 2018). Removing the Ca^2+^-independent signal produced IC catalogs similar to the catalogs produced from the uncorrected signal. Furthermore, the functional connectivity analyses on hemodynamic corrected signals revealed connectivity changes across behavior periods similar to those found with uncorrected signals. Therefore, the additional dual wavelength results agree with previous reports that the hemodynamic contribution to GCaMP6 Ca^2+^ response is limited (Vanni and Murphy 2014; Makino et al. 2017; Allen et al. 2017; Jacobs et al. 2018; Musall et al. 2019).

### Decrease in connectivity of primary motor and somatosensory regions

Primary motor and somatosensory nodes decrease in correlations, both at the onset and throughout locomotion. Furthermore, when compared to rest, the centrality of somatosensory nodes markedly decreases during locomotion. It would have been reasonable to expect an increase connectivity in these nodes, since M1 modulates gait, which inherently requires receiving upstream sensory inputs of those obstacles (Drew and Marigold 2015). However, our results suggest that premotor regions may send sensory-to-motor translation calculations to M1 and other cortical regions, as discussed below.

Lesioning the corticospinal tract or M1 produces locomotor deficits, including hypermetria and abnormalities in limb trajectory and intralimb coordination (Liddell and Phillips 1944). Pyramidal neuronal firing in M1 correlates with individual muscle activity during locomotion (Drew et al. 2008a; Drew et al. 2008b; Drew and Marigold 2015) and modulates when the subject maneuvers around an obstacle, suggesting M1 directly controls locomotion and influences the central motor pattern. The observed decrease in correlations of M1 nodes potentially reflects the role of M1 in accurate limb placement, a function not known to be found elsewhere in the cerebral cortex. Interestingly, this decrease in connectivity begins before the onset of locomotion, which agrees with the long-known finding that M1 is activated during movement preparation (Tanji and Evarts 1976; Georgopoulos et al. 1989; Churchland et al. 2010).

### Increase in connectivity of retrosplenial regions

In contrast to primary motor cortical nodes, retrosplenial nodes display increased functional connectivity with the majority of the dorsal cerebral cortex. These increases in connectivity occur when comparing rest with preparation and rest with continued locomotion and agree with previous findings showing increased retrosplenial connectivity during locomotion (Clancy et al. 2019). This corroborates previous evidence that the retrosplenial region is approximately equivalent to the posterior parietal cortex (PPC) in carnivores and primates and is engaged in integrating the sensory and spatial information needed to accurately and safely navigate the environment during locomotion (Drew and Marigold 2015; Takakusaki 2017). For example, animals with lesioned PPCs are unable to retain information of obstacles once the obstacle passes out of the visual field (Lajoie et al. 2010). The largest increase occurs before the onset of locomotion, as would be expected if the retrosplenial region is tasked with assessing the environment for navigational routes or potential obstacles at the start of locomotion. As there was no change in the magnitude or direction of Granger causality, the ratio of information being sent and received by retrosplenial nodes does not change from rest to locomotion.

### Increase in connectivity and outward causality of anterior M2 regions

Our third and perhaps most notable finding is the increase in functional connectivity of the most anterior nodes of M2. The increase first occurs at the onset of locomotion and remains elevated through the offset. These anterior M2 nodes increase in correlation with somatosensory, parietal, visual, and retrosplenial regions during locomotion and show a significant increase in outward Granger causality to these regions. While we acknowledge that Granger causality is not a direct measure of cause-and-effect, it is a valuable tool for estimating directed connectivity between brain regions (Seth et al. 2015; Barnett et al. 2018). Our results corroborate the observed increases in correlation between V1 and M2 during locomotion in the mouse (Clancy et al. 2019) and suggests the more regions of the premotor cortex provide an organizing signal to the rest of the cerebral cortex during locomotion. This agrees with previous studies that show the premotor cortex modulates individual primary sensory regions (Schneider et al. 2014; Nelson and Mooney 2016; Leinweber et al. 2017).

The function of the premotor cortex in locomotion is not well understood (Drew and Marigold 2015), but it is known to be essential in several discrete motor behaviors. In mice, these behaviors include licking (Chen et al. 2017; Allen et al. 2017; Inagaki et al. 2018) and lever pressing(Makino et al. 2017). In such tasks, M2 activity can be used to causally predict activity in other dorsal cortical regions during movement (Makino et al. 2017), and inactivation of M2 represses cortex-wide responses to sensory stimuli (Allen et al. 2017), suggesting M2 has widespread influence over the activity of the cerebral cortex. Furthermore, premotor cortex is widely anatomically connected, receiving inputs from the somatosensory, auditory, posterior parietal, and orbital cortices, and projecting to primary motor, somatosensory, parietal, and retrosplenial areas (Yamawaki et al. 2016; Zhang et al. 2016; Leinweber et al. 2017; Lin et al. 2018).

What process is the premotor cortex carrying out through this widespread connectivity and influence over the cortex? In the primate, the premotor cortex is involved in transforming sensory information for movement planning in discrete movements (Di Pellegrino G. and Ladavas 2015). Multimodal neurons in the ventral premotor cortex map visual and auditory stimuli in spatial relation to the space immediately surrounding the body, or “peri-personal space” (Rizzolatti et al. 1981a; Rizzolatti et al. 1981b; Graziano et al. 1994; Fogassi et al. 1996). Peri-personal space is important for both motor and cognitive computations (Serino 2019), and inhibiting the premotor cortex impairs reaction times to stimuli in peri-personal space (Serino et al. 2011). Stimuli within peri-personal space modulate the motor system, including decreasing the excitability of M1 (Makin et al. 2009; Serino et al. 2009), and the premotor cortex is necessary for this modulation (Avenanti et al. 2012). During locomotion, the representation of physical distance from the body expands during walking, even when other sensory cues are stationary (Noel et al. 2015). Moreover, locomotive behavior is facilitated when salient stimuli appear far away rather than up close (Di Marco S. et al. 2019). These effects likely improve obstacle avoidance during locomotion (Noel et al. 2015) and/or promote movement toward a salient stimulus (Di Marco S. et al. 2019).

Therefore, the evidence above suggests that M2 provides an interface between sensory and motor systems to create a model of the outside world that can be used by M1 to execute appropriate motor responses and by sensory regions to properly interpret incoming stimuli. In support of this hypothesis, the present results show a significant increase in connectivity and causality from the anterior M2 to motor, sensory, and integrative regions across the dorsal cerebral cortex. The findings suggest anterior M2 drives cortical locomotion computation by sending an organizing signal to the rest of the cerebral cortex.

## Acknowledgements

We would like to thank Lijuan Zhuo for her assistance in animal surgeries. The Minnesota Supercomputing Institute provided high-processing computing, and the University Imaging Centers provided 3D printing services of the cortical implants. This work was supported by National Institutes of Health (NS111028 to S.B.K. and T.J. E., R61/R33 NS115089 to T.J.E., and T32-MH115688 to S.L.W., as well as by a grant from University of Minnesota Informatics Institute, which includes support from the University of Minnesota’s MnDRIVE Initiative (to S.L.W).

---

**Supplemental Figure 1.**
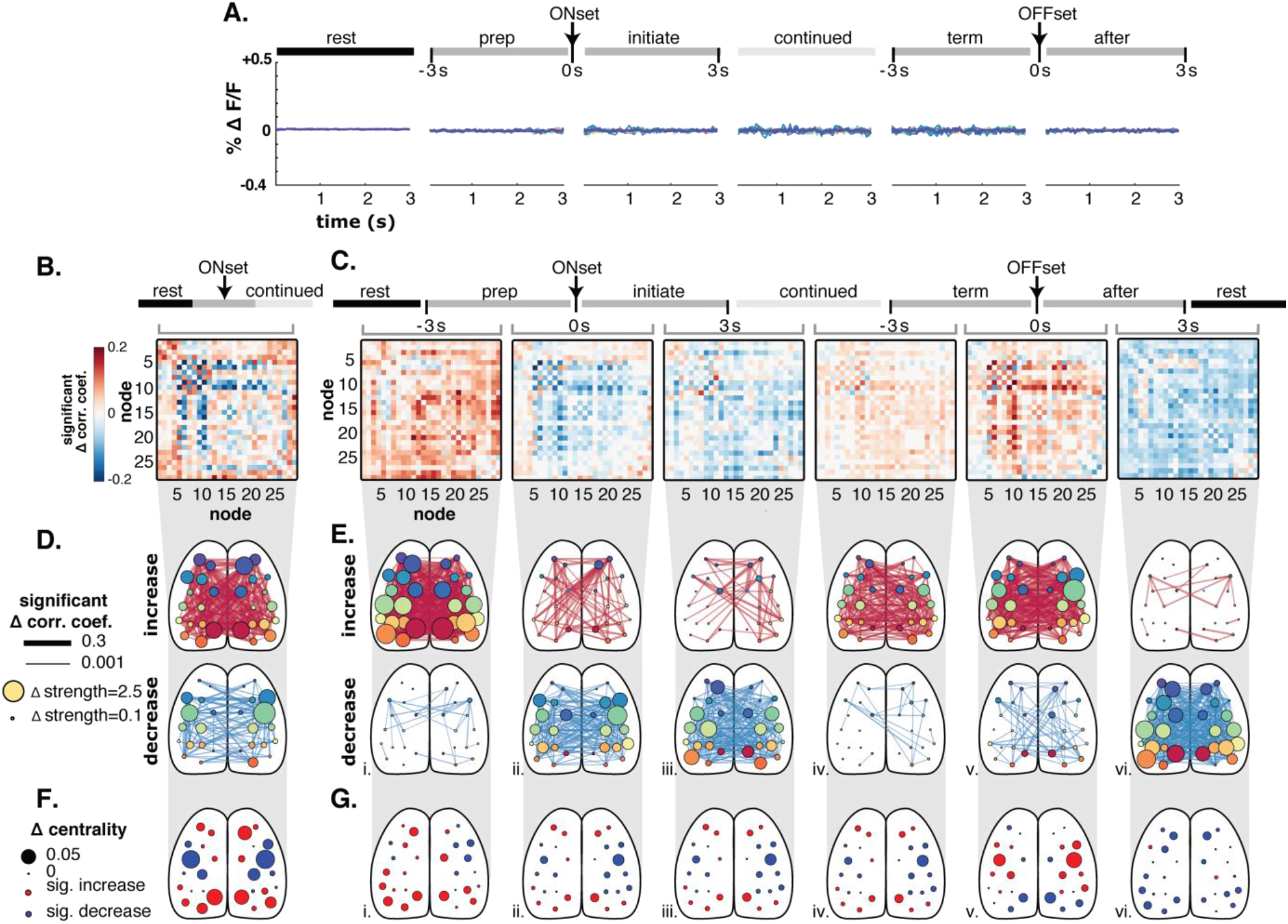
High pass filtering does not alter measures of functional connectivity. **A.** Removal of fluorescence signal below 2 Hz removes the average increases in fluorescence observed with locomotion (see Fig. 3C). Results in B-G are based on high pass filtered data. **B** and **C.** Adjacency matrix representation of significant increases and decreases from rest to continued locomotion (B) and across behavior periods (C). **D** and **E.** Significant increases and decreases from rest to continued locomotion (D) and across behavior periods (E) shown in graph representations superimposed on the cortical surface. **F** and **G.** Adjacency matrix representation of significant changes in eigenvector centrality from rest to continued locomotion (F) and across behavior periods (G). Trends in functional connectivity found in unfiltered data are preserved across behavior periods (see Fig. 5B and C) as are trends in eigenvector centrality (see Fig. 6B and C).

**Supplemental Figure 2.**
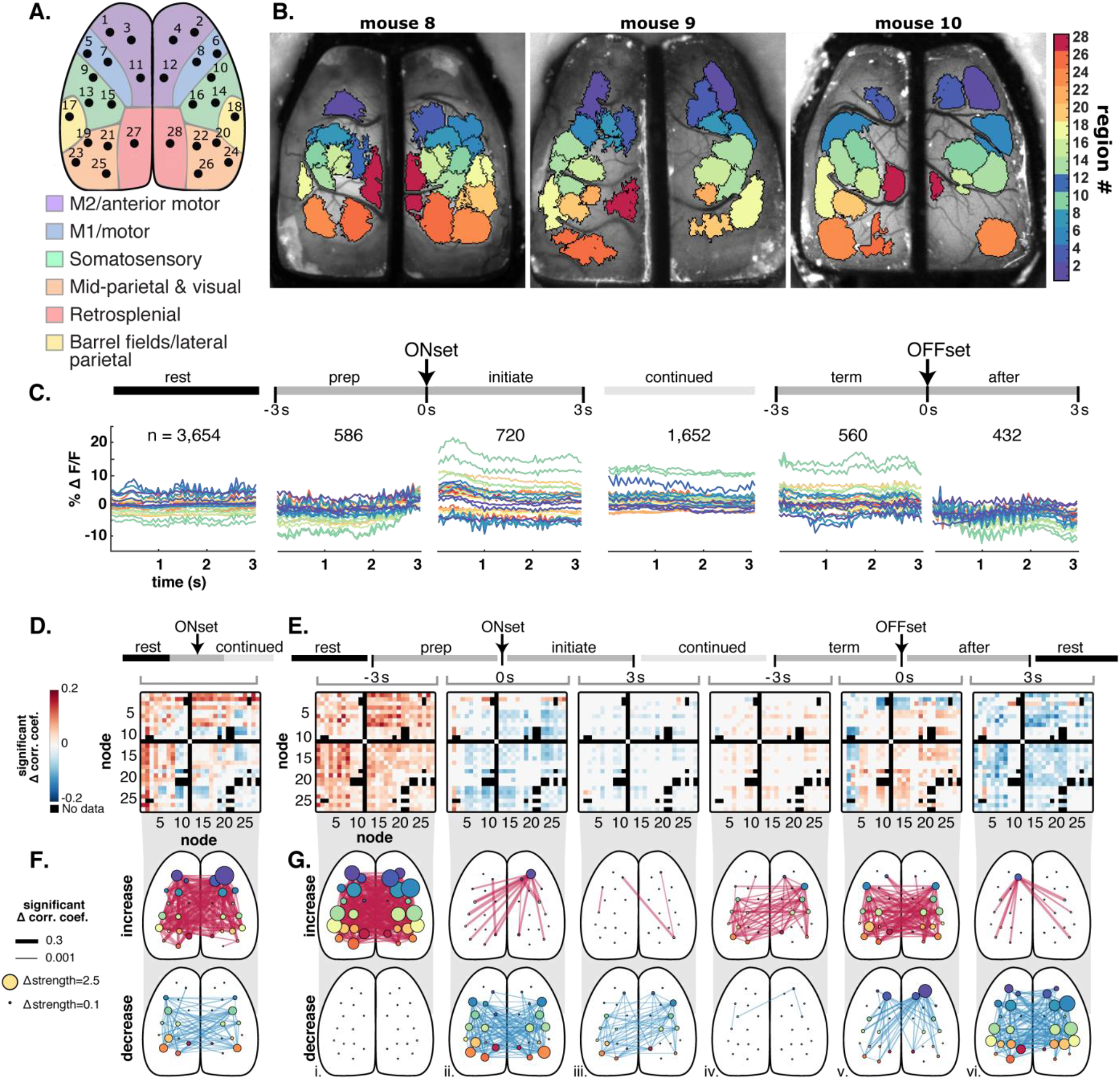
Results of hemodynamic corrected imaging data. **A.** The 28 nodes found to be consistent across mice in the main analysis were also used to group the ICs found in the hemodynamic corrected data. Node 11 did not occur in any of these 3 mice. **B.** The resulting IC catalogs for each of the 3 mice included in the hemodynamic correction data and the corresponding node assignments. **C**. Fluorescence activity from the common set of 28 nodes averaged over all mice and across all instances of each 3s behavior period (n=number of periods averaged). Abbreviations as in Figure 3. **D.** Matrix of significant changes in correlation between nodes from rest to continued locomotion (α<0.05, permutation test with false discovery rate correction). **E** Matrix of significant changes in correlations across sequential behavior periods, calculated and displayed as in **D. F.** Significant increases (top) and decreases (bottom) from rest to continued locomotion shown in graph representations, as in Figure. 5. **G**. Significant increases (top) and decreases (bottom) in correlations across sequential behavior periods, as in F.

## References

1. Allen Institute for Brain Science. 2015. Technical white paper: Allen mouse common coordinate framework. Allen Institute for Brain Science. 1:1–18.

2. Allen WE, Kauvar IV, Chen MZ, Richman EB, Yang SJ, Chan K, Gradinaru V, Deverman BE, Luo L, Deisseroth K. 2017. Global representations of goal-directed behavior in distinct cell types of mouse neocortex. Neuron. 94(4):891–907.

3. Avenanti A, Annela L, Serino A. 2012. Suppression of premotor cortex disrupts motor coding of peripersonal space. Neuroimage. 63(1):281–8.

4. Ayaz A, Stauble A, Hamada M, Wulf MA, Saleem AB, Helmchen F. 2019. Layer-specific integration of locomotion and sensory information in mouse barrel cortex. Nat Commun. 10(1):2585.

5. Barnett L, Barrett AB, Seth AK. 2018. Solved problems for Granger causality in neuroscience: a response to Stokes and Purdon. Neuroimage. 178:744–8.

6. Barnett L, Seth AK. 2014. The MVGC multivariate Granger causality toolbox: a new approach to Granger-causal inference. J Neurosci Methods. 223:50–68.

7. Buzsaki G. 2010. Neural syntax: cell assemblies, synapsembles, and readers. Neuron. 68(3):362–85.

8. Cardoso JF. 1999. High-order contrasts for independent component analysis. Neural Comput. 11(1):157–92.

9. Carrillo-Reid L, Yang W, Bando Y, Peterka DS, Yuste R. 2016. Imprinting and recalling cortical ensembles. Science. 353(6300):691–4.

10. Chapin JK, Woodward DJ. 1982. Somatic sensory transmission to the cortex during movement: phasic modulation over the locomotor step cycle. Exp Neurol. 78(3):670–84.

11. Chen TW, Li N, Daie K, Svoboda K. 2017. A map of anticipatory activity in mouse motor cortex. Neuron. 94(4):866–79.

12. Churchland MM, Cunningham JP, Kaufman MT, Ryu SI, Shenoy KV. 2010. Cortical preparatory activity: representation of movement or first cog in a dynamical machine? Neuron. 68(3):387–400.

13. Clancy KB, Orsolic I, Mrsic-Flogel TD. 2019. Locomotion-dependent remapping of distributed cortical networks. Nat Neurosci. 22(5):778–786.

14. Dadarlat MC, Stryker MP. 2017. Locomotion enhances neural encoding of visual stimuli in mouse V1. J Neurosci. 37(14):3764–75.

15. Dana H, Chen TW, Hu A, Shields BC, Guo C, Looger LL, Kim DS, Svoboda K. 2014. Thy1-GCaMP6 transgenic mice for neuronal population imaging in vivo. PLoS ONE. 9(9):e108697.

16. Di Marco S., Tosoni A, Altomare EC, Ferretti G, Perrucci MG, Committeri G. 2019. Walking-related locomotion is facilitated by the perception of distant targets in the extrapersonal space. Sci Rep. 9(1):9884.

17. Di Pellegrino G., Ladavas E. 2015. Peripersonal space in the brain. Neuropsychologia. 66:126–33.

18. Dipoppa M, Ranson A, Krumin M, Pachitariu M, Carandini M, Harris KD. 2018. Vision and locomotion shape the interactions between neuron types in mouse visual cortex. Neuron. 98(3):602–15.

19. Drew T, Andujar JE, Lajoie K, Yakovenko S. 2008a. Cortical mechanisms involved in visuomotor coordination during precision walking. Brain Res Rev. 57(1): 199–211.

20. Drew T, Kalaska J, Krouchev N. 2008b. Muscle synergies during locomotion in the cat: a model for motor cortex control. J Physiol. 586(5): 1239–45.

21. Drew T, Marigold DS. 2015. Taking the next step: cortical contributions to the control of locomotion. Curr Opin Neurobiol. 33:25–33.

22. Economo MN, Viswanathan S, Tasic B, Bas E, Winnubst J, Menon V, Graybuck LT, Nguyen TN, Smith KA, Yao Z, Wang L, Gerfen CR, Chandrashekar J, Zeng H, Looger LL, Svoboda K. 2018. Distinct descending motor cortex pathways and their roles in movement. Nature. 563(7729):79–84.

23. Favorov OV, Nilaweera WU, Miasnikov AA, Beloozerova IN. 2015. Activity of somatosensory-responsive neurons in high subdivisions of SI cortex during locomotion. J Neurosci. 35(20):7763–76.

24. Ferezou I, Haiss F, Gentet LJ, Aronoff R, Weber B, Petersen CC. 2007. Spatiotemporal dynamics of cortical sensorimotor integration in behaving mice. Neuron. 56(5):907–23.

25. Fogassi L, Gallese V, Fadiga L, Luppino G, Matelli M, Rizzolatti G. 1996. Coding of peripersonal space in inferior premotor cortex (area F4). J Neurophysiol. 76(1): 141–57.

26. Freedman DJ, Ibos G. 2018. An Integrative Framework for Sensory, Motor, and Cognitive Functions of the Posterior Parietal Cortex. Neuron. 97(6): 1219–34.

27. Genovese CR, Lazar NA, Nichols T. 2002. Thresholding of statistical maps in functional neuroimaging using the false discovery rate. Neuroimage. 15(4):870–8.

28. Georgopoulos AP, Crutcher MD, Schwartz AB. 1989. Cognitive spatial-motor processes. 3. Motor cortical prediction of movement direction during an instructed delay period. Exp Brain Res. 75(1):183–94.

29. Ghanbari L, Carter RE, Rynes M, Dominguez J, Chen G, Naik A, Hu J, Sagar MAK, Halton L, Mossazaghi N, Gray MM, West SL, Eliceiri KW, Ebner TJ, Kodandaramaiah SB. 2019. Cortex-wide neural interfacing via transparent polymer skulls. Nature Commun. 10(1): 1500.

30. Ghose GM. 2015. Vision and vigilance on the go. Trends Cogn Sci. 19(3): 115–6.

31. Graziano MS, Yap GS, Gross CG. 1994. Coding of visual space by premotor neurons. Science. 266(5187):1054–7.

32. Guizar M. 2021. Efficient subpixel image registration by cross-correlation. MATLAB central file exchange. (https://www.mathworks.com/matlabcentral/fileexchange/18401-efficient-subpixel-image-registration-by-cross-correlation) Retrieved March 5, 2021.

33. Han Y, Kebschull JM, Campbell RAA, Cowan D, Imhof F, Zador AM, Mrsic-Flogel TD. 2018. The logic of single-cell projections from visual cortex. Nature. 556(7699):51–6.

34. Harris KD. 2005. Neural signatures of cell assembly organization. Nat Rev Neurosci. 6(5):399–407.

35. Hernandez A, Nacher V, Luna R, Zainos A, Lemus L, Alvarez M, Vazquez Y, Camarillo L, Romo R. 2010. Decoding a perceptual decision process across cortex. Neuron. 66(2):300–14.

36. Hira R, Ohkubo F, Tanaka YR, Masamizu Y, Augustine GJ, Kasai H, Matsuzaki M. 2013. In vivo optogenetic tracing of functional corticocortical connections between motor forelimb areas. Front Neural Circuits. 7:55.

37. Inagaki HK, Inagaki M, Romani S, Svoboda K. 2018. Low-dimensional and monotonic preparatory activity in mouse anterior lateral motor cortex. J Neurosci. 38(17):4163–85.

38. Jacobs EAK, Steinmetz NA, Carandini M, Harris KD. 2018. Cortical state fluctuations during sensory decision making. Curr Biol. 30(24):4944–4955.

39. Josset N, Roussel M, Lemieux M, Lafrance-Zoubga D, Rastqar A, Bretzner F. 2018. Distinct contributions of mesencephalic locomotor region nuclei to locomotor control in the freely behaving mouse. Curr Biol. 28(6):884–901.

40. Kauvar IV, Machado TA, Yuen E, Kochalka J, Choi M, Allen WE, Wetzstein G, Deisseroth K. 2020. Cortical observation by synchronous multifocal optical sampling reveals widespread population encoding of actions. Neuron. 107(2):351–67.

41. Keller GB, Bonhoeffer T, Hubener M. 2012. Sensorimotor mismatch signals in primary visual cortex of the behaving mouse. Neuron. 74(5):809–15.

42. Lajoie K, Andujar JE, Pearson K, Drew T. 2010. Neurons in area 5 of the posterior parietal cortex in the cat contribute to interlimb coordination during visually guided locomotion: a role in working memory. J Neurophysiol. 103(4):2234–54.

43. Lee AM, Hoy JL, Bonci A, Wilbrecht L, Stryker MP, Niell CM. 2014. Identification of a brainstem circuit regulating visual cortical state in parallel with locomotion. Neuron. 83(2):455–66.

44. Lee KY, Li M, Manchanda M, Batra R, Charizanis K, Mohan A, Warren SA, Chamberlain CM, Finn D, Hong H, Ashraf H, Kasahara H, Ranum LP, Swanson MS. 2013. Compound loss of muscleblind-like function in myotonic dystrophy. EMBO Mol Med. 5(12):1887–900.

45. Leinweber M, Ward DR, Sobczak JM, Attinger A, Keller GB. 2017. A sensorimotor circuit in mouse cortex for visual flow predictions. Neuron. 96(5): 1204.

46. Li J. 2015. Reconstructing neuronal connectivity from calcium imaging data using generalized transfer entropy. Public Health Theses. (Yale Univ) 1179.

47. Liddell EGT, Phillips CG. 1944. Pyramidal section in the cat. Brain. 67(1):1–9.

48. Lin HM, Kuang JX, Sun P, Li N, Lv X, Zhang YH. 2018. Reconstruction of intratelencephalic neurons in the mouse secondary motor cortex reveals the diverse projection patterns of single neurons. Front Neuroanat. 12:86.

49. Ma Y, Shaik MA, Kim SH, Kozberg MG, Thibodeaux DN, Zhao HT, Yu H, Hillman EM. 2016. Wide-field optical mapping of neural activity and brain haemodynamics: considerations and novel approaches. Philos Trans R Soc Lond B Biol Sci. 371(1705).

50. MacDowell CJ, Buschman TJ. 2020. Low-Dimensional Spatio-Temporal Dynamics Underlie Cortex-Wide Neural Activity. Curr Biol. 30(14):2665–2680.

51. Makin TR, Holmes NP, Brozzoli C, Rossetti Y, Farne A. 2009. Coding of visual space during motor preparation: Approaching objects rapidly modulate corticospinal excitability in hand-centered coordinates. J Neurosci. 29(38):11841–51.

52. Makino H, Ren C, Liu H, Kim AN, Kondapaneni N, Liu X, Kuzum D, Komiyama T. 2017. Transformation of cortex-wide emergent properties during motor learning. Neuron. 94(4):880–890.

53. Mao D, Molina LA, Bonin V, McNaughton BL. 2020. Vision and Locomotion Combine to Drive Path Integration Sequences in Mouse Retrosplenial Cortex. Curr Biol. 30(9):1680–1688.

54. McGinley MJ, David SV, McCormick DA. 2015. Cortical Membrane Potential Signature of Optimal States for Sensory Signal Detection. Neuron. 87(1):179–192.

55. Mohajerani MH, Chan AW, Mohsenvand M, Ledue J, Liu R, McVea DA, Boyd JD, Wang YT, Reimers M, Murphy TH. 2013. Spontaneous cortical activity alternates between motifs defined by regional axonal projections. Nat Neurosci. 16(10):1426–35.

56. Musall S, Kaufman MT, Juavinett AL, Gluf S, Churchland AK. 2019. Single-trial neural dynamics are dominated by richly varied movements. Nat Neurosci. 22(10): 1677–86.

57. Neafsey EJ, Sievert C. 1982. A second forelimb motor area exists in rat frontal cortex. Brain Res. 232(1):151–6.

58. Nelson A, Mooney R. 2016. The basal forebrain and motor cortex provide convergent yet distinct movement-related inputs to the auditory cortex. Neuron. 90(3):635–48.

59. Niell CM, Stryker MP. 2010. Modulation of visual responses by behavioral state in mouse visual cortex. Neuron. 65(4):472–9.

60. Noel JP, Grivaz P, Marmaroli P, Lissek H, Blanke O, Serino A. 2015. Full body action remapping of peripersonal space: the case of walking. Neuropsychologia. 70:375–84.

61. Parker PRL, Brown MA, Smear MC, Niell CM. 2020. Movement-related signals in sensory areas: roles in natural behavior. Trends Neurosci. 43(8):581–95.

62. Petersen TH, Willerslev-Olsen M, Conway BA, Nielsen JB. 2012. The motor cortex drives the muscles during walking in human subjects. J Physiol. 590(10):2443–52.

63. Pinto L, Rajan K, DePasquale B, Thiberge SY, Tank DW, Brody CD. 2019. Task-dependent changes in the large-scale dynamics and necessity of cortical regions. Neuron. 104(4):810–24.

64. Polack PO, Friedman J, Golshani P. 2013. Cellular mechanisms of brain state-dependent gain modulation in visual cortex. Nat Neurosci. 16(9): 1331–9.

65. Rizzolatti G, Scandolara C, Matelli M, Gentilucci M. 1981a. Afferent properties of periarcuate neurons in macaque monkeys. I. Somatosensory responses. Behav Brain Res. 2(2): 125–46.

66. Rizzolatti G, Scandolara C, Matelli M, Gentilucci M. 1981b. Afferent properties of periarcuate neurons in macaque monkeys. II. Visual responses. Behav Brain Res. 2(2): 147–63.

67. Rubinov M, Sporns O. 2010. Complex network measures of brain connectivity: uses and interpretations. Neuroimage. 52(3):1059–69.

68. Sahonero-Alvarez G, Calderon H. 2017. A comparison of SOBI, FastICA, JADE and infomax algorithms. IMCIC 8th International Multi-Conference on Complexity, Informatics and Cybernetics, Proceedings.

69. Saleem AB, Ayaz A, Jeffery KJ, Harris KD, Carandini M. 2013. Integration of visual motion and locomotion in mouse visual cortex. Nat Neurosci. 16(12):1864–9.

70. Schneider DM. 2020. Reflections of action in sensory cortex. Curr Opin Neurobiol. 64:53–9.

71. Schneider DM, Mooney R. 2018. How movement modulates hearing. Annu Rev Neurosci. 41:553–72.

72. Schneider DM, Nelson A, Mooney R. 2014. A synaptic and circuit basis for corollary discharge in the auditory cortex. Nature. 513(7517): 189–94.

73. Serino A. 2019. Peripersonal space (PPS) as a multisensory interface between the individual and the environment, defining the space of the self. Neurosci Biobehav Rev. 99:138–59.

74. Serino A, Annella L, Avenanti A. 2009. Motor properties of peripersonal space in humans. PLoS ONE. 4(8):e6582.

75. Serino A, Canzoneri E, Avenanti A. 2011. Fronto-parietal areas necessary for a multisensory representation of peripersonal space in humans: an rTMS study. J Cogn Neurosci. 23(10):2956–67.

76. Seth AK, Barrett AB, Barnett L. 2015. Granger causality analysis in neuroscience and neuroimaging. J Neurosci. 35(8):3293–7.

77. Sharma S, Kim LH, Whelan PJ. 2019. Towards a connectome of descending commands controlling locomotion. Curr Opin in Physiol. 8:70–75

78. Steinmetz NA, Zatka-Haas P, Carandini M, Harris KD. 2019. Distributed coding of choice, action and engagement across the mouse brain. Nature. 576(7786):266–273.

79. Takakusaki K. 2017. Functional neuroanatomy for posture and gait control. J Mov Disord. 10(1):1–17.

80. Tang L, Higley MJ. 2020. Layer 5 circuits in V1 differentially control visuomotor behavior. Neuron. 105(2):346–54.

81. Tanji J, Evarts EV. 1976. Anticipatory activity of motor cortex neurons in relation to direction of an intended movement. J Neurophysiol. 39(5): 1062–8.

82. Vanni MP, Chan AW, Balbi M, Silasi G, Murphy TH. 2017. Mesoscale mapping of mouse cortex reveals frequency-dependent cycling between distinct macroscale functional modules. J Neurosci. 37(31):7513–33.

83. Vanni MP, Murphy TH. 2014. Mesoscale transcranial spontaneous activity mapping in GCaMP3 transgenic mice reveals extensive reciprocal connections between areas of somatomotor cortex. J Neurosci. 34(48):15931–46.

84. Xiao D, Vanni MP, Mitelut CC, Chan AW, LeDue JM, Xie Y, Chen AC, Swindale NV, Murphy TH. 2017. Mapping cortical mesoscopic networks of single spiking cortical or sub-cortical neurons. Elife. 6:e19976.

85. Yamawaki N, Radulovic J, Shepherd GM. 2016. A corticocortical circuit directly links retrosplenial cortex to M2 in the mouse. J Neurosci. 36(36):9365–74.

86. Yang J-H, Kwan AC. 2020. Secondary motor cortex: broadcasting and biasing animal’s decisions through long-range circuits. International Rev of Neurobiol. (in press).

87. Zagha E, Casale AE, Sachdev RN, McGinley MJ, McCormick DA. 2013. Motor cortex feedback influences sensory processing by modulating network state. Neuron. 79(3):567–78.

88. Zhang S, Xu M, Chang WC, Ma C, Hoang Do JP, Jeong D, Lei T, Fan JL, Dan Y. 2016. Organization of long-range inputs and outputs of frontal cortex for top-down control. Nat Neurosci. 19(12): 1733–42.

89. Zhou M, Liang F, Xiong XR, Li L, Li H, Xiao Z, Tao HW, Zhang LI. 2014. Scaling down of balanced excitation and inhibition by active behavioral states in auditory cortex. Nat Neurosci. 17(6):841–50.

90. Zingg B, Hintiryan H, Gou L, Song MY, Bay M, Bienkowski MS, Foster NN, Yamashita S, Bowman I, Toga AW, Dong HW. 2014. Neural networks of the mouse neocortex. Cell. 156(5):1096–111.

